# Interleukin-6 restricts pre-thymic T cell lineage commitment of progenitors driving loss of SIV control

**DOI:** 10.64898/2026.01.29.702123

**Authors:** Saleem Anwar, Naseem Sadek, Christian Michel Beusch, Ahmet F. Coskun, Mohamed S. Abdel-Hakeem, R. Paul Johnson, Frank J. T. Staal, Vijayakumar Velu, Mirko Paiardini, Brandon F. Keele, Guido Silvestri, David Ezra Gordon, Jeffrey A. Tomalka, Sheikh Abdul Rahman

## Abstract

Effective T cell reconstitution in people living with HIV is central to durable immune control and cure strategies. Sustained thymic output underpins T cell recovery and requires continuous seeding by T cell-committed progenitors originating in the bone marrow (BM). Using the SIV/rhesus macaque model, we identified a thymus-seeding progenitor (TSP; CD4⁻CD8⁻CD34⁺CD38⁻CD7⁺) in BM declining rapidly following SIV infection. This loss closely associated with reduction in T cell lineage committed differentiation of BM-derived hematopoietic stem and progenitor cells (HSPCs). Importantly, both the decline in TSPs and the impairment of pre-thymic T cell potential were strongly associated with early loss of viral control, independent of peripheral T cell dynamics. Plasma interleukin-6 (IL-6) levels robustly predicted the magnitude of TSP loss and the restriction of T cell-biased HSPC differentiation. Integrated transcriptomic and proteomic analyses revealed inflammatory imprinting of HSPCs characterized by activation of the IL-6-JAK-STAT axis, inflammasome engagement, and coordinated suppression of key T cell specification factors, including RUNX1, FYN, and ZAP70. In a nonanimal model of thymopoiesis, IL-6 exposure of rhesus macaque and human HSPCs inhibited their transition from DN1 (CD38⁻) to DN2 (CD38⁺) TSP states, indicating an early block in T cell lineage commitment. Conversely, ex vivo IL-6 receptor blockade restored thymocyte differentiation to levels comparable to untreated controls. Collectively, these findings demonstrate that pathogenic inflammation restricts pre-thymic T cell development early after infection, directly contributing to loss of viral control. These findings have important implications for understanding the mediators of anti-viral T cell immunity and HIV cure.

## Introduction

Chronic immune activation represents a central paradox of HIV/AIDS pathogenesis: while the host mounts a sustained inflammatory response, this activation fails to mediate viral clearance and instead drives disease progression(1, 2). Despite suppressive antiretroviral therapy (ART), people living with HIV (PLWH) exhibit persistent inflammation that predicts both AIDS and non-AIDS morbidity(3, 4). A critical consequence of this chronic inflammatory state is the impairment of thymic function, which limits the regenerative capacity of the T cell compartment(5–12). Understanding the mechanisms linking systemic inflammation to defective T cell regeneration is essential for developing strategies to restore immune competence in HIV infection.

Sustained thymopoiesis depends on the continuous recruitment of bone marrow (BM)-derived hematopoietic stem and progenitor cells (HSPCs) to the thymus(13, 14). These thymus-seeding progenitors (TSPs), phenotypically characterized as CD4^−^CD8^−^ double-negative (DN) CD34^+^CD7^+^ cells, colonize the thymus and initiate T cell development(15–17). Upon thymic entry, TSPs undergo a conserved developmental program: upregulation of CD38 marks the transition from the DN1 (CD34^+^CD38^−^CD7^+^) to DN2 (CD34^+^CD38^+^CD7^+^) stage, followed by CD34 downregulation, emergence of CD4^+^CD3− immature single-positive (ISP) thymocytes, progression through the CD4^+^CD8^+^ double-positive (DP) intermediate, and ultimate maturation into single-positive CD4^+^ or CD8^+^ T cells(17–21). Thus, the T cell lineage commitment of HSPCs in the BM represents a critical pre-thymic checkpoint for maintaining thymic seeding and peripheral T cell homeostasis.

HSPC fate decisions are dynamically regulated by microenvironmental cues, with inflammatory signals exerting profound effects on lineage output(22–34). Under conditions of acute or chronic inflammation, HSPCs preferentially adopt a myeloid lineage bias through emergency myelopoiesis, a process that enhances innate immune cell production at the expense of lymphopoiesis(30, 33–35). This inflammation-driven skewing of hematopoiesis has direct relevance to HIV infection, where T cells are essential for controlling viremia. Notably, rare individuals termed elite controllers maintain viral suppression without ART and exhibit characteristically low levels of inflammation(36, 37). Conversely, immunological non-responders—individuals who fail to recover CD4^+^ T cell counts despite viral suppression on therapy—display elevated inflammatory markers(38–41). These clinical observations suggest that the inflammatory milieu may fundamentally alter the T cell regenerative capacity of BM-derived HSPCs.

Here, we tested the hypothesis that pathogenic inflammation impairs the T cell lineage specification of BM-derived HSPCs directly contributing to viral control. Using the SIV/rhesus macaque model of HIV/AIDS, combined with thymus independent artificial thymic organoid systems for both macaque (RhATO; (21)) and human (hATO;(19)) progenitors, we demonstrate that BM-derived HSPCs rapidly lose their capacity to generate T cell-lineage committed progenitors following SIV infection directly translating to pre-thymic T cell regenerative deficiency of HSPCs independent of thymus dysfunction. This decline in T cell regenerative potential correlates with early loss of viral control. We identify interleukin-6 (IL-6) as a key inflammatory mediator that directly inhibits T cell specification of HSPCs by suppressing the DN1-to-DN2 transition during thymopoiesis. Importantly, IL-6 receptor blockade reverses this inhibitory effect. These findings demonstrate IL-6-mediated inflammatory signaling as a previously unrecognized mechanism under lentiviral infection directly suppressing T cell lineage commitment, and this pre-thymic paucity of T cell regenerative potential directly translating into loss of viral control, very early post infection – independent of peripheral T cell dynamics.

## Results

### T cell committed progenitors in bone marrow declined early upon pathogenic SIV infection

To characterize thymus-seeding progenitors (TSPs) in bone marrow (BM) of SIV-infected rhesus macaques (RMs), we analyzed CD4⁻CD8⁻ double-negative (DN) CD34⁺CD7⁺ cells, including the CD38⁻ TSP subset previously shown to exhibit T cell lineage-biased differentiation in humans(17, 42), in matched BM and thymus samples from eight SIV-naïve adult RMs (Fig. 1a–i; Supplementary Fig. 1a–j). DN CD34⁺CD7⁺ TSPs constituted approximately 3.2% (geomean) of total DN CD34⁺ HSPCs (∼914 cells/10⁶ mononuclear cells; geomean) in BM compared with ∼35% (geomean; ∼871 cells/10⁶ thymocytes) in thymus (Fig. 1d, e). Although the thymus exhibited >10-fold higher TSP proportions relative to BM, absolute TSP numbers per 10⁶ cells were comparable between compartments (Fig. 1e). The frequency of CD7-expressing DN CD34⁺ HSPCs showed significant positive correlation between BM and thymus (r = 0.80, P < 0.01; Fig. 1f), which was lost upon exclusion of the CD7 marker (r = −0.30, P > 0.05; Fig. 1g). Furthermore, BM-derived CD38⁻CD34⁺CD7⁺ frequency (analogous to human DN1 TSPs(17, 42)) exhibited strong associations with thymic progenitor populations and T-cell factor-1 (TCF-1)-expressing progenitors (Fig. 1h, i). Functional validation using our rhesus artificial thymic organoid (RhATO) system (21) confirmed T cell developmental ontogeny, generating CD4⁺CD3⁻ immature single-positive (ISP), CD4⁺CD8⁺ double-positive (DP), and mature CD3⁺ single-positive T cell subsets from BM-derived DN CD34⁺CD7⁺ TSPs (Fig. 1j–l; Supplementary Fig. 2a–c).

**Figure 1.**
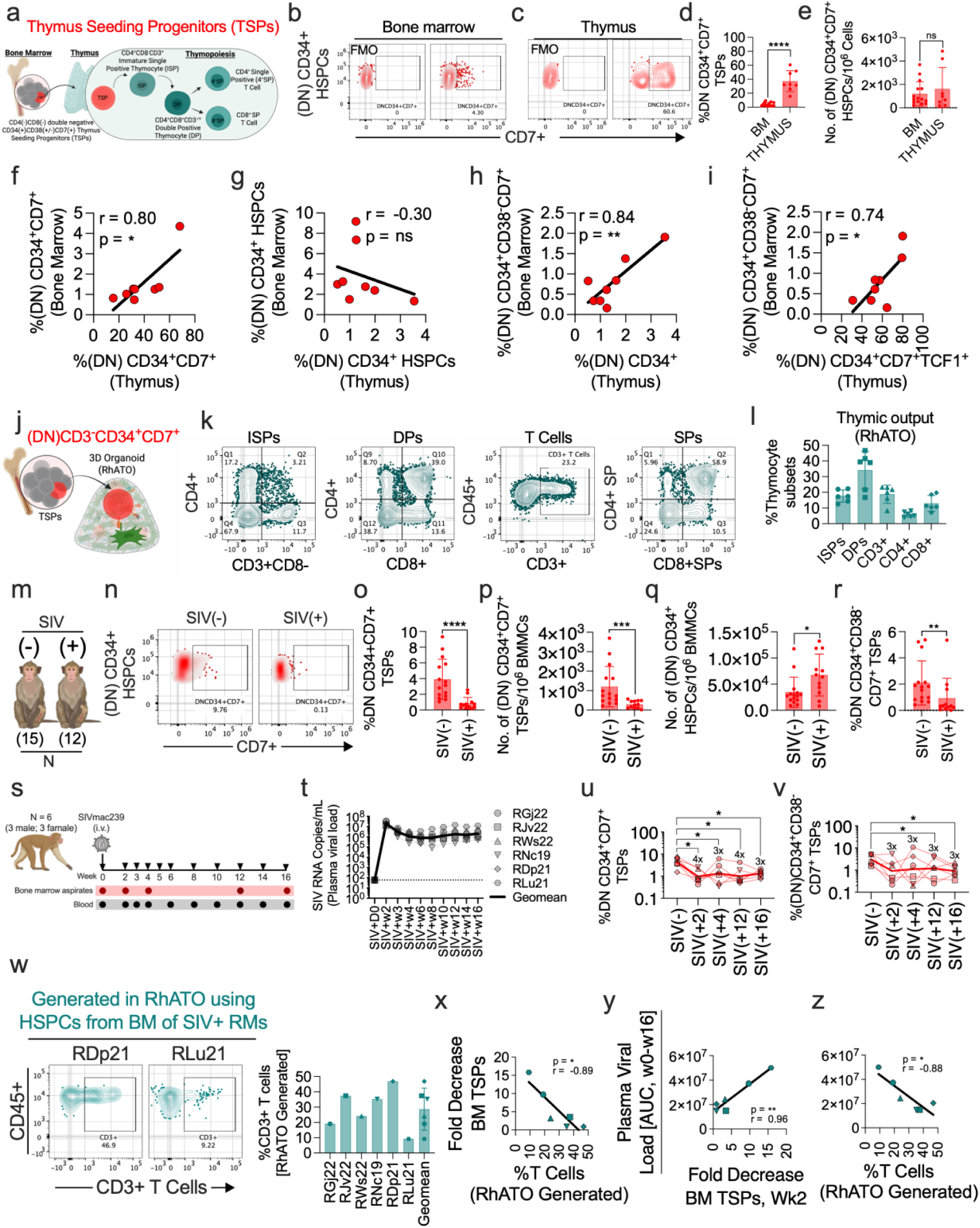
Loss of T cell committed progenitors in bone marrow under SIV infection drive loss of viral control (a-z). **a,** Schematic illustrating thymus-seeding progenitors (TSPs) arising from double-negative (DN; CD4⁻CD8⁻) CD34⁺CD7⁺ hematopoietic stem and progenitor cells (HSPCs) in the bone marrow and their migration to the thymus, where they undergo stepwise differentiation through CD4⁺CD3^−^ immature single-positive (ISP), CD4⁺CD8⁺ double-positive (DP), and single-positive (SP) CD4⁺ or CD8⁺ T-cell stages. **b-c,** Representative flow-cytometry plots showing (DN)CD34⁺CD7⁺ HSPCs in bone marrow (b) and thymus (c), with corresponding fluorescence-minus-one (FMO) controls. **d,** Frequency of (DN)CD34⁺CD7⁺ TSPs among bone-marrow and thymic HSPCs. **e,** Absolute numbers of (DN)CD34⁺CD7⁺ HSPCs per 10^6^ total bone marrow mononuclear cells and thymocytes analyzed. **f–i,** Pearson correlation analyses between matched (N=8) bone-marrow and thymus derived CD4^−^CD8^−^ double negative CD34^+^ progenitor populations, comparing frequencies of CD7 expressing (f), and non-expressing (g) DN CD34⁺ HSPCs. **h,** Pearson correlation between bone marrow derived CD34⁺CD38^−^CD7⁺ (DN1) TSPs and thymus derived (DN)CD34+ (h), and (DN)CD34+CD7+TCF1+ (i) progenitors. Pearson correlation coefficients (r) and P values are shown. **j,** Schematic of the previously described Rhesus-specific nonanimal model of thymopoiesis known as Rhesus-artificial thymic organoid (RhATO) system (21), seeded with CD3^−^(DN)CD34⁺CD7⁺ TSPs. **k,** Representative flow-cytometry plots illustrating T cell lineage committed differentiation intermediates representing frequencies of ISPs, and DPs early thymocyte intermediates, mature CD3⁺ T cells, as well as CD4⁺ and CD8⁺ single positive T cells, analyzed at week 4 of organoid establishment as previously described (k). **l,** Bar graph showing quantification of the frequencies of indicated thymocyte subsets generated in the RhATO-system (N=6 RhATOs). **m,** Schematic depicting uninfected (SIV⁻; N=15) and SIV-infected (SIV⁺; N=12) rhesus macaques (RMs) used to analyze bone marrow derived progenitors. **n,** Representative flow-cytometry plots showing (DN)CD34⁺CD7⁺ T cell committed progenitors in bone marrow of SIV⁻ and SIV⁺ RMs. **o,** Bar graph showing quantitation of the frequency of (DN)CD34⁺CD7⁺ TSPs among bone-marrow HSPCs in SIV⁻ and SIV⁺ RMs. **p,** Absolute numbers of (DN)CD34⁺CD7⁺ T-cell committed progenitors per 10⁶ bone-marrow mononuclear cells (BMMCs) analyzed. **q,** Absolute numbers of (DN)CD34⁺ HSPCs per 10⁶ BMMCs analyzed. **r,** Frequency of CD34⁺CD38^−^CD7⁺ (DN1) TSPs among bone-marrow (DN)CD34+CD7+ progenitors in SIV⁻ and SIV⁺ animals. **s,** Experimental design showing RMs (N=6) infected with 300 TCID_50_ SIVmac239M via intravenous route. Bone marrow aspirates (red dots) and blood (black dots) collected at the indicated timepoints starting with week 0 of infection. **t,** Line graph showing longitudinal plasma viral load (PVL) at the indicated timepoints. Geomean PVL indicated in black and individual data indicated in grey. **u-v,** Line graphs showing longitudinal frequencies of (DN)CD34⁺CD7⁺ (u) and CD34⁺CD38^−^CD7⁺ (DN1) TSPs (v) TSPs at baseline (week 0) and at week 2, 4, 12, and 16 following SIV infection. **w,** Representative flow-cytometry plots and bar graph showing frequencies of T cells generated in RhATO cultures using 5000 CD3^−^CD34^+^ HSPCs per RhATO isolated from bone marrow of RMs at week 16 of infection. **x,** Pearson correlation analysis between fold decrease of CD34^+^CD38^−^CD7^+^ (DN1) TSP population at week 2 of infection and frequency of T cells generated by CD3^−^CD34^+^ HSPCs in RhATO culture. **y–z,** Pearson correlation analyses between total plasma viral burden until week 16 post infection represented as area under curve and fold decrease of CD34^+^CD38^−^CD7^+^ (DN1) TSP (y), and frequency of T cells generated in RhATO (z). Each symbol represents one animal. Statistical significance was determined using Mann Whitney unpaired two-tailed t-tests for unrelated data sets, Pearson correlation, or paired Wilcoxon test for related data sets; (*P < 0.05; **P < 0.01; ***P < 0.001; ****P < 0.0001).

Next, we analyzed the impact of SIV infection on frequency of TSPs in BM (Fig. 1m–r). SIV-infected RMs (n = 12; various stages of chronic infection) demonstrated >5-fold lower DN CD34⁺CD7⁺ TSP frequency, declining from 3.2% (n = 15; SIV-naïve) to 0.58% (P < 0.001; Fig. 1n–p), with a corresponding 4-fold decline in absolute cell numbers per 10⁶ BM mononuclear cells (Fig. 1p). Notably, total DN CD34⁺ HSPC numbers increased ∼2-fold under SIV infection (Fig. 1q), while the frequency of CD38⁻ (DN1) TSP subset also demonstrated ∼5-fold reduction in bone marrow of SIV infected RMs (Fig. 1r). Longitudinal sampling in six RMs at day 0 and weeks 2, 4, 12, and 16 post-infection (p.i.) revealed significant decline (>4-fold) in DN CD34⁺CD7⁺ TSP frequency as early as week 2 p.i., coinciding with peak viremia, with partial recovery at week 4 followed by subsequent decline at week 16 (Fig. 1s–v). Similarly, the DN CD34⁺CD38⁻CD7⁺ (DN1 TSP) subset decreased ∼3-fold at week 2 p.i. and remained significantly reduced through week 16 (Fig. 1v). These results identify the CD34⁺CD38⁻CD7⁺ (DN1) TSPs in bone marrow of RMs, analogous to humans, and demonstrate rapid depletion of T cell-committed progenitors in bone marrow as early as week 2 p.i. despite increased in total HSPC numbers in SIV infected RMs.

### Restricted pre-thymic T cell lineage commitment of bone marrow progenitors associated with early loss of viral control independent of peripheral T cells

To study a possible impact of reduced T cell lineage commitment of bone marrow progenitors on de novo generation of T cells, bone marrow CD3⁻CD34⁺ HSPCs were FACS-sorted from six RMs at week 16 p.i. and differentiated into T cells as previously described(21). A 5-fold difference in frequencies of ex vivo-generated T cell was observed (∼25% geometric mean; range: 9–47%; Fig. 1w; Supplementary Fig. 2d), whereas frequencies of peripheral blood T cells at week 16 p.i. showed a distinct dynamic (geomean frequency ∼34%; range: 25.6–41.2%; Supplementary Fig. 2e, f). Notably, the magnitude of CD34⁺CD38⁻CD7⁺ (DN1) TSPs decline observed in bone marrow of RMs at week 2 p.i. correlated significantly with proportionate decreases in T cell differentiation potential (r = 0.89, P < 0.01; Fig. 1x), and both parameters associated with reduced peripheral T cell frequencies in 4/6 RMs at week 16 p.i. (Supplementary Fig. 3a, b).

Correlation analysis revealed total viral load (AUC; weeks 0–16 p.i.) strongly associating with both, reduced CD34⁺CD38⁻CD7⁺ (DN1) TSP frequency in bone marrow at week 2 and impaired T cell differentiation potential obtained in RhATO cultures (Fig. 1y, z). Further, fold drop of (DN1) TSP frequencies observed at week 2p.i. was strongly associated with loss of viral control at all subsequent time points post infection (Supplementary Fig. 3c–f), as was decreased pre-thymic T cell differentiation potential (Supplementary Fig. 2g–j). Further, 4/6 RMs also showed strong association between loss of viral control and decrease in peripheral CD3+ T cells (Supplementary Fig. 3k) independent of pre-thymic ex-vivo generated T cells directly from bone marrow CD3^−^CD34^+^ HSPCs. These results suggest that early TSP decline directly primes T cell developmental deficits driving loss of viral control during chronic lentiviral infection.

### Plasma IL-6 level associated with reduced T cell lineage commitment of bone marrow progenitors

To investigate associations between plasma viral load (PVL), systemic inflammation, and T cell commitment of BM progenitors, we measured plasma cytokines and chemokines in six longitudinally tracked RMs at day 0 and weeks 2, 4, and 16 p.i. (Fig. 2). PVL peaked at week 2 p.i., showing 2 × 10⁷ SIV RNA copies/mL plasma (geometric mean), declined to 1.12 × 10⁶ copies/mL at week 4, and rebounded to 2.16 × 10⁶ copies/mL at week 16 (Fig. 2a). Plasma IL-6 exhibited parallel cyclical dynamics with 5-fold increase at week 2 (P = 0.015), 5-fold decrease at week 4 (P = 0.015), and 2.5-fold increase at week 16 (P = 0.046; Fig. 2b). Other pro-inflammatory cytokines (IL-15, TNF-β, IL-1β) increased following infection but exhibited distinct temporal dynamics (Fig. 2c–e; Supplementary Fig. 4a–d). Of all the chemokines analyzed, interferon gamma inducible protein (IP-10)/CXCL10, a ligand for CXCR3 receptor expressed on HSPCs and implicated in reducing their self-renewal capacity as well as their myeloid skewing(43), reported significant (3-fold; P=0.015) and persistent increase through week 16 (Fig. 2f; Supplementary Fig. 4e–h).

**Figure 2.**
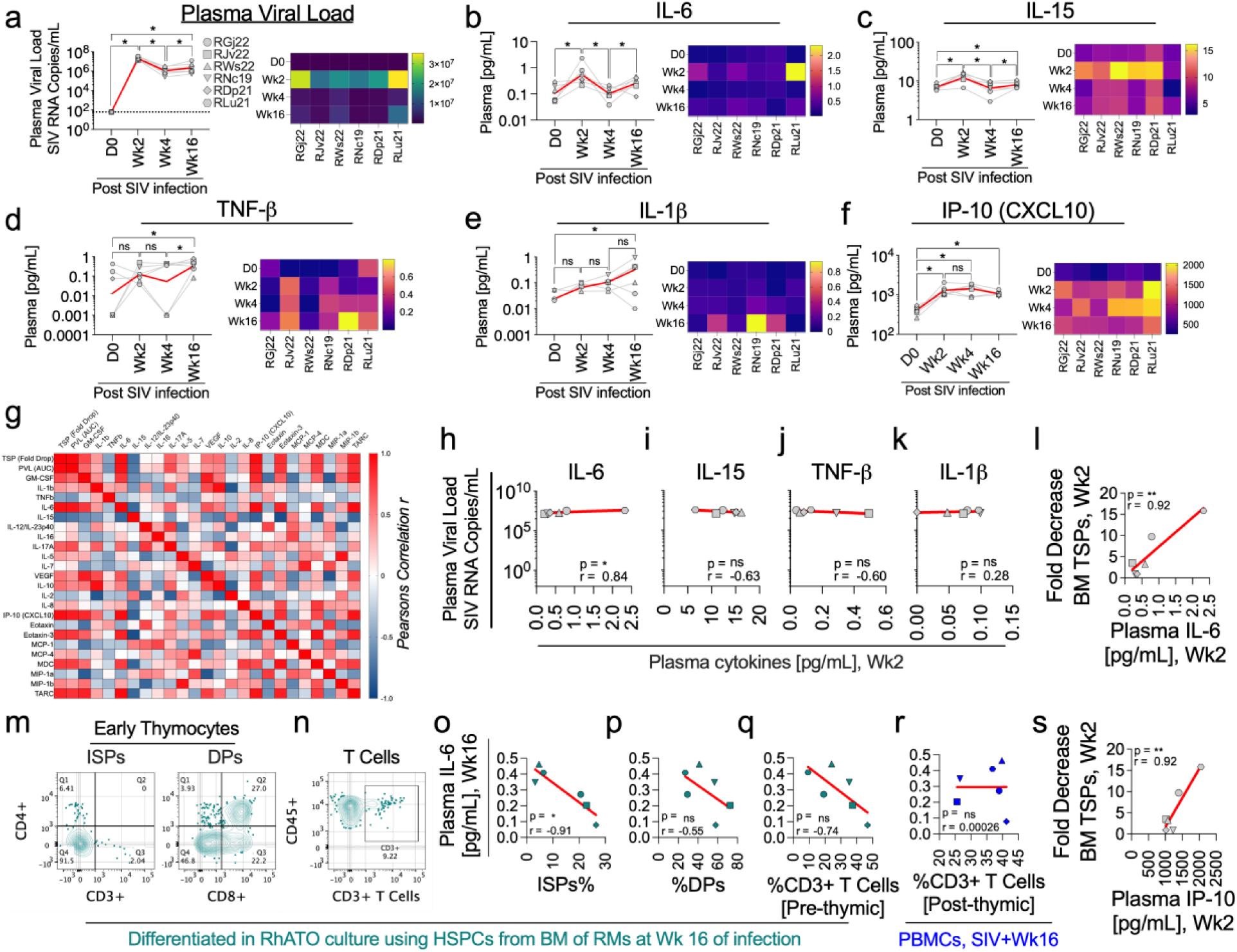
Plasma IL-6 associated strongly with reduced T cell committed progenitors and differentiation (a-s). **a**, Longitudinal plasma viral load (PVL) in SIV-infected rhesus macaques (RMs) measured before infection (D0) and at week (Wk) 2, 4 and 16 post SIV infection (N=6). **b–f**, Line graphs and corresponding heatmaps representing longitudinal plasma concentrations of IL-6 (**b**), IL-15 (**c**), TNF-β (**d**) IL-1β (**e**) and IP-10 (CXCL10) (**f**) across the same time points. (**g)**, Pearson correlation matrices summarizing associations between CD34⁺CD38^−^CD7⁺ (DN1) thymic seeding progenitors (TSPs) attrition, plasma viral load and circulating cytokines and chemokines during peak viremia (week 2 post SIV). Pearson correlation coefficients (r) is shown. (**h–k)**, Pearson correlation between peak PVL and corresponding plasma concentrations of IL-6 (**h**), IL-15 (**i**), TNF-β (**j**) IL-1β (**k**) at week 2 post infection. **l,** Pearson correlation between (DN1) TSP attrition and plasma IL-6 concentrations at peak viremia (week 2) post infection. **(m-n),** Representative flow-cytometry plots illustrating RhATO-derived T cell developmental ontogeny from bone marrow CD3^−^CD34^+^ HSPCs of RMs isolated at week 16 of infection, culture analyzed at week 4 of establishment. Flow plots show differentiation stages of thymocytes including CD4^+^CD3^−^ immature single positive (ISP) early thymocyte, CD4^+^CD8^+^ double positive (DP) intermediate thymocyte, (m) and mature CD3⁺ T cells (n). **o-q,** Pearson correlation analyses between plasma IL-6 at week 16 of infection and RhATO derived frequencies of ISPs (o), DPs (p), and CD3+ T cells (q) stages. **r,** Pearson correlation analyses between plasma IL-6 at week 16 of infection and frequency of CD3+ T cells in blood of RMs at week 16 post infection. **s**, Pearson correlation between fold decrease in frequency of bone marrow (DN1) TSPs and plasma IP-10 concentrations. Each symbol represents one animal; lines indicate longitudinal measurements from the same animal. Heat-map color scales indicate relative concentrations or viral load. Linear regression lines, Pearson correlation coefficients (r) and P value calculated by paired Wilcoxon test are shown where appropriate; (*P < 0.05; **P < 0.01).

Correlation analysis revealed IL-6 as the only cytokine significantly associated with PVL (r = 0.93, P = 0.007; Fig. 2g–h). No correlation between PVL and IL-15 (Fig. 2g,i), TNF-β (Fig. 2g,j), or IL-1β (Fig. 2g,k) was observed. At peak viremia, plasma IL-6 levels correlated significantly with TSP decline (r = 0.92, P = 0.008; Fig. 2l), whereas lower IL-15 levels associated with higher TSP decline (Supplementary Fig. 4i). At week 16 p.i., plasma IL-6 associated with decreased T cell lineage-committed differentiation, with the strongest correlation observed at the CD4⁺CD3⁻ ISP early thymocyte stage (r = −0.91, P < 0.01; Fig. 2m–q; Supplementary Fig. 4l–o). Notably, IL-6 did not associate with the frequencies of peripheral T cells analyzed at week 16 p.i. (Fig. 2r). Plasma IP-10 at peak viremia (week 2) significantly associated with decreased BM TSPs (r = −0.92, P < 0.009; Fig. 2s). These results identify IL-6 and IP-10 as early predictors of impaired T cell commitment in BM progenitors, suggesting early pre-thymic restriction in T cell lineage commitment of bone marrow progenitors.

### Early loss of T cell-committed progenitors in the bone marrow paralleled their impaired T cell lineage differentiation capacity

To confirm restricted T cell lineage committed differentiation of bone marrow progenitors under chronic lentiviral infection, we analyzed 31 RhATOs generated from 15 SIV-naïve RMs and 34 RhATOs from 12 SIV-infected RMs (various stages of chronic infection), assessing progression to early thymic differentiation stage of culture before any potential homeostatic proliferation of distinct developmental stages confound a reliable estimation of thymic restriction (Fig. 3a–f). We have shown that at week 1 of RhATO culture most prominent early thymic intermediate observed was CD4+CD3-ISPs(21). At week 1 of RhATO culture, frequency of CD4⁺CD3⁻ ISP early thymocyte intermediates from SIV-infected RM-derived HSPCs (∼6% geometric mean) showed 4-fold decrease (P = 0.0001) compared with SIV-naïve RMs (∼24% geometric mean), despite comparable CD45⁺ lymphocyte differentiation between groups (∼68% vs. ∼64%; Fig. 3e). The ISP-to-lymphocyte ratio confirmed significant 4-fold reduction (P = 0.0001) in thymocyte differentiation potential (Fig. 3f).

**Figure 3.**
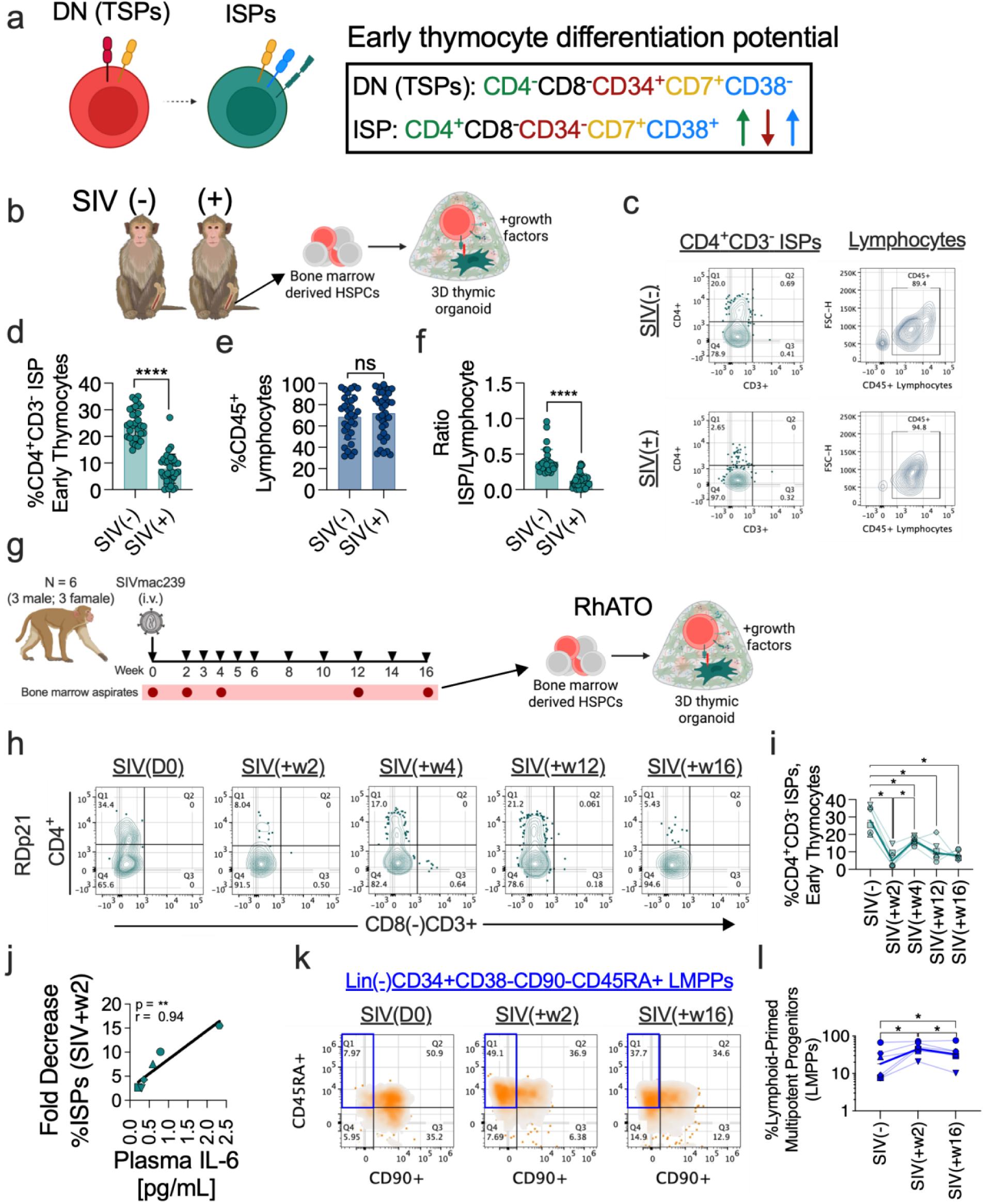
Bone-marrow derived CD34^+^CD3^−^ HSPCs exhibited early decline in their T cell lineage committed differentiation potential (a-l). **a,** Cartoon illustrating early T cell lineage committed differentiation steps from CD4⁻CD8⁻CD34⁺CD38⁻CD7⁺ (DN1) TSPs to CD4^+^CD3^−^ immature single-positive (ISPs) thymocyte stage, highlighting early phenotypic changes during T cell committed differentiation process. **b,** Experimental schematic illustrating isolation of CD3^−^CD34^+^ hematopoietic stem and progenitor cells (HSPCs) from bone-marrow of SIV-uninfected (SIV⁻; N = 15) and SIV-infected (SIV⁺; N = 12) rhesus macaques (RMs) and their differentiation to ISP thymocyte in the rhesus-artificial thymic organoid (RhATO) system. **c,** Representative flow-cytometric plots showing frequencies of ISPs and corresponding CD45⁺ lymphocytes generated in RhATO culture at week 1 of culture as previously described(21). **d,** Bar graph showing quantitation of the frequency of ISP thymocytes for SIV⁻ (N=31 RhATOs) and SIV⁺ (N=34 RhATOs) conditions. **e,** Bar graph showing corresponding frequency of CD45⁺ lymphocytes. **f,** Bar graph showing ratio of the frequencies of ISP thymocytes to CD45⁺ lymphocytes generated in RhATO system. **g,** Schematic showing longitudinal ex-vivo tracking of the T cell lineage committed differentiation potential of the CD3^−^CD34^+^ HSPCs isolated from bone marrow of six RMs at the indicated timepoints (red dots) post SIV infection. **h,** Representative flow-cytometric plot showing frequencies of the ISP thymocytes generated in RhATOs. Each plot represents each timepoint post infection T cell lineage committed potential of HSPCs were evaluated. **i,** Line graph showing geomean (dark green line) and individual data points (lighter green lines) of the frequencies of CD4⁺CD3⁻ ISP thymocytes generated in RhATOs from bone-marrow HSPCs collected at the indicated time points following SIV infection. **j,** Pearson correlation between drop in T cell lineage committed differentiation potential of bone marrow derived HSPCs (as estimated by frequency of ex-vivo generated ISPs in RhATO cultures at week 2 compared with day 0 post infection) and plasma IL-6 concentrations at week 2 post infection. **k,** Representative flow-cytometric plots identifying Lin⁻CD34⁺CD38⁻CD90⁻CD45RA⁺ lymphoid-primed multipotent progenitors (LMPPs) in bone marrow at baseline (SIV D0), week 2 and week 16 post-infection. **l,** Line graph showing geomean (dark blue) and individual values (lighter blue lines) frequencies of LMPPs in bone marrow at day 0, week 2 and 16 post infection. Each symbol represents an individual animal. Data are presented as geomean ± geomean standard deviation. Statistical significance was determined using Mann Whitney unpaired two-tailed t-tests for unrelated data sets, Pearson correlation, or paired Wilcoxon test for related data sets; (*P < 0.05; ****P < 0.0001; ns, not significant).

Longitudinal analysis of CD3⁻CD34⁺ HSPCs isolated from bone marrow of RMs at day 0 and weeks 2, 4, 12, and 16 p.i. revealed ∼6-fold decrease (P = 0.015) in their ISP differentiation potential (∼5% vs. ∼28% at day 0; Fig. 3h-i; Supplementary Fig. 5a, b) at week 2p.i. T-cell lineage committed differentiation showed 4-fold recovery (P = 0.015) at week 4 compared with week 2, followed by gradual decline through week 16. Plasma IL-6 at week 2 strongly correlated with decreased thymocyte differentiation potential (r = 0.94, P = 0.008; Fig. 3j). Phenotypic analysis of lymphoid-primed multipotent progenitors (LMPPs; Lin⁻CD34⁺CD38⁻CD90⁻CD45RA⁺) based on reported gating strategy (31) revealed >2-fold increase (P = 0.015) in their frequency from ∼18% (day 0) to ∼46% (week 2), declining modestly to ∼32% at week 16 (Fig. 3k, l; Supplementary Fig. 5c). These results demonstrate cyclic decline of T cell differentiation concomitant with TSP depletion, while lymphoid progenitor differentiation remains uninterrupted.

### Bone marrow HSPCs cycled between lineage- uncommitted and committed state under SIV infection

To investigate impact of persistent lentiviral infection on bone marrow CD34+ HSPC differentiation priming states, we analyzed lineage marker expression (CD2, CD3, CD7, CD10, CD11b, CD19, CD20, CD33, CD56, CD235a) in BM from 15 SIV-naïve and 12 SIV-infected RMs (Fig. 4a, b). Frequency of Lin⁻CD34⁺ HSPCs increased 2-fold (P = 0.01) under SIV-infection (∼5% geometric mean) compared with no infection control (∼2%). Longitudinal tracking in six RMs revealed ∼4-fold increase (P = 0.015) in proportion of Lin⁻CD34⁺ HSPCs at week 2 p.i., ∼2-fold decrease (P = 0.04) at week 4, followed by cyclical fluctuations at weeks 12 and 16, remaining significantly elevated above pre-infection levels throughout (Fig. 4c).

**Figure 4.**
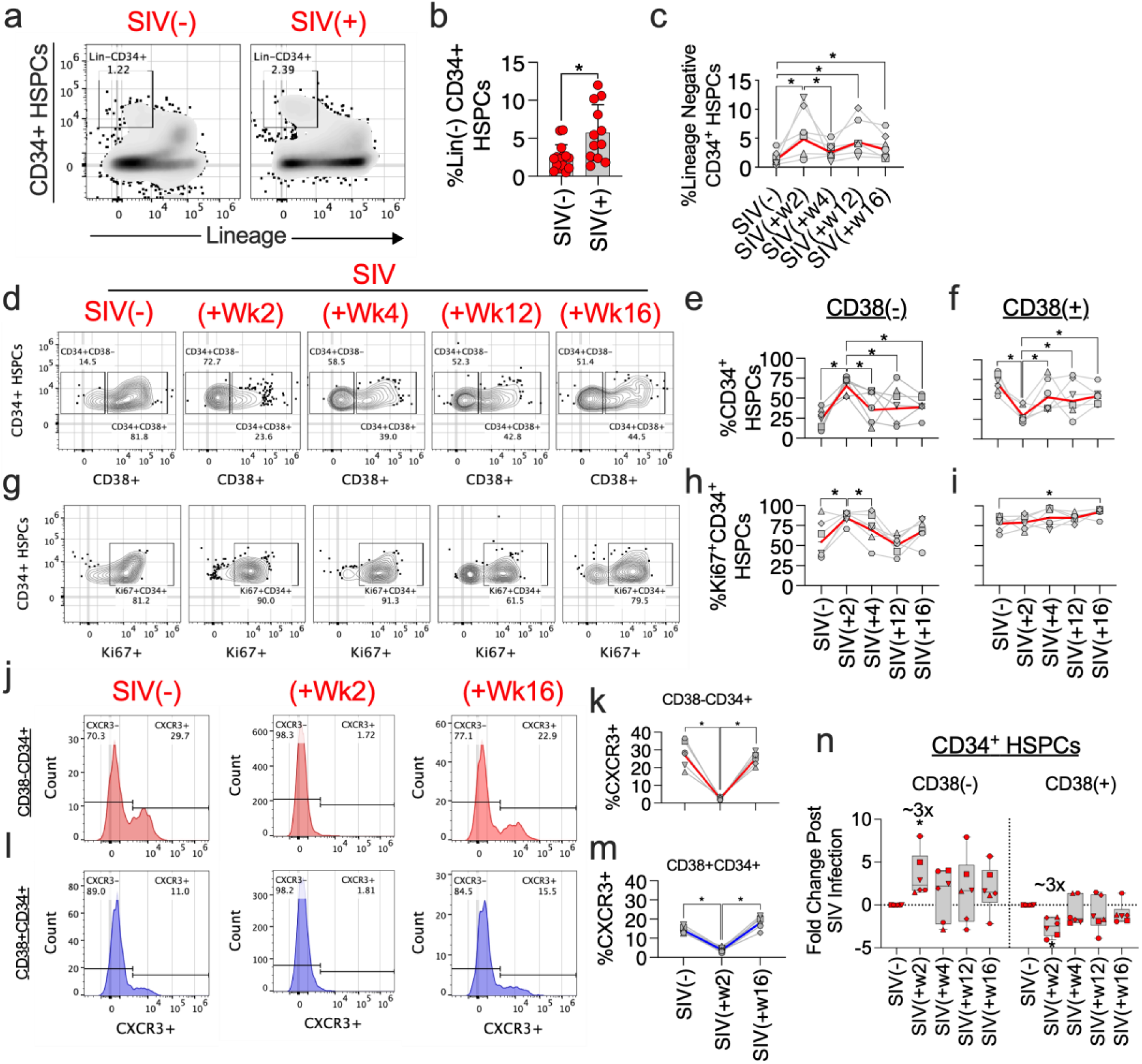
Bone marrow derived HSPCs demonstrated a cyclic shift between undifferentiated and differentiated state under pathogenic SIV infection (a-n). **a**, Representative flow-cytometry plots showing lineage-negative (Lin⁻) CD34⁺ hematopoietic stem and progenitor cells (HSPCs) in SIV⁻ (N=15) and SIV⁺ (N=12) Rhesus macaques (RMs). Numbers indicate frequencies within the indicated gates. **b**, Frequency of Lin⁻ CD34⁺ HSPCs in SIV⁻ and SIV⁺ RMs. **c**, Longitudinal analysis of Lin⁻ CD34⁺ HSPC frequencies at day 0 (SIV^−^) and at week 2, 4, 12 and 16 post SIV infection in six RMs. **d**, Representative flow-cytometry plots showing subdivision of CD34⁺ HSPCs into CD38⁻ and CD38⁺ subsets before infection (SIV⁻) and at weeks 2, 4, 12 and 16 post SIV infection. **e-f**, Longitudinal frequencies of CD38⁻ (**e**) and CD38⁺ (**f**) cells within the CD34⁺ HSPC compartment across indicated time points post infection. **g**, Representative flow-cytometry plots depicting Ki67 expression within total CD34⁺ HSPCs at baseline and following SIV infection. **h-i**, Frequencies of Ki67⁺ cells within CD34⁺CD38⁻ (**h**) and CD34⁺CD38⁺ (**i**) HSPC subsets across the indicated time points. **J-k**, Representative histograms showing CXCR3 surface expression on CD34⁺CD38⁻ (**j**, red) and Longitudinal quantitation of the frequency of CXCR3⁺CD34⁺CD38- (**k**, red) HSPCs before infection (SIV⁻) and at weeks 2 and 16 post SIV infection. **l-m**, Representative histograms showing CXCR3 surface expression on CD34⁺CD38⁺ (**l**, blue) and Longitudinal quantitation of the frequency of CXCR3⁺CD34⁺CD38⁺ (**m**, red) HSPCs before infection (SIV⁻) and at weeks 2 and 16 post SIV infection. **n**, Fold change relative to pre-infection baseline in CD34⁺ HSPC subsets following SIV infection, analyzed separately for CD38⁻ and CD38⁺ compartments. Each symbol represents one RM; lines connect longitudinal samples from the same animal. Red symbols and lines denote group means. Statistical significance was determined using Mann Whitney unpaired two-tailed t-tests for unrelated data sets, or paired Wilcoxon t-tests for related data sets; *P < 0.05).

Next, we analyzed expression of CD38 on CD34^+^ HSPCs, which distinguished primitive/dormant (CD38⁻) from committed (CD38⁺) HSPC states (Fig. 4d–f). Longitudinal analysis post infection revealed 3-fold increase in the frequency of CD38⁻CD34⁺ HSPCs (P = 0.015) from ∼23% (day 0) to ∼65% (week 2), decreasing to ∼35% at week 4 and ∼38% at week 16. Concomitantly, frequency of CD38⁺CD34⁺ HSPCs decreased >2-fold (P = 0.015) from ∼70% (day 0) to ∼30% (week 2), partially recovering to ∼52% at week 4 but remaining significantly reduced through week 16 (P < 0.05). The CD38⁻CD34⁺ fraction, but not CD38⁺CD34⁺, exhibited significantly higher Ki-67 expression (P = 0.015; ∼85%) at week 2, declining to ∼69% at week 4 (Fig. 4g–i). CXCR3 expression, a key receptor for IP-10 (CXCL10) ligand, showed overall significantly higher expression on CD38^−^ subset of HSPCs (Supplementary Fig. 6a-c). Frequency of CXCR3^+^CD38⁻CD34⁺ HSPCs declined >10-fold at week 2 before returning to baseline at week 16 p.i. (Fig. 4j, k). Though lower compared to CD38⁻CD34⁺, expression on CD38^+^CD34⁺ also showed similar dynamics (Fig. 4l-m). Overall, CD34⁺ HSPCs at week 2 exhibited maximal shift toward a Lin⁻CXCR3⁻Ki-67⁺CD38⁻ phenotype (Fig. 5n), suggesting HSPCs acquire a more primitive, undifferentiated state early upon infection, followed by cyclical transitions between differentiated and undifferentiated states.

**Figure 5.**
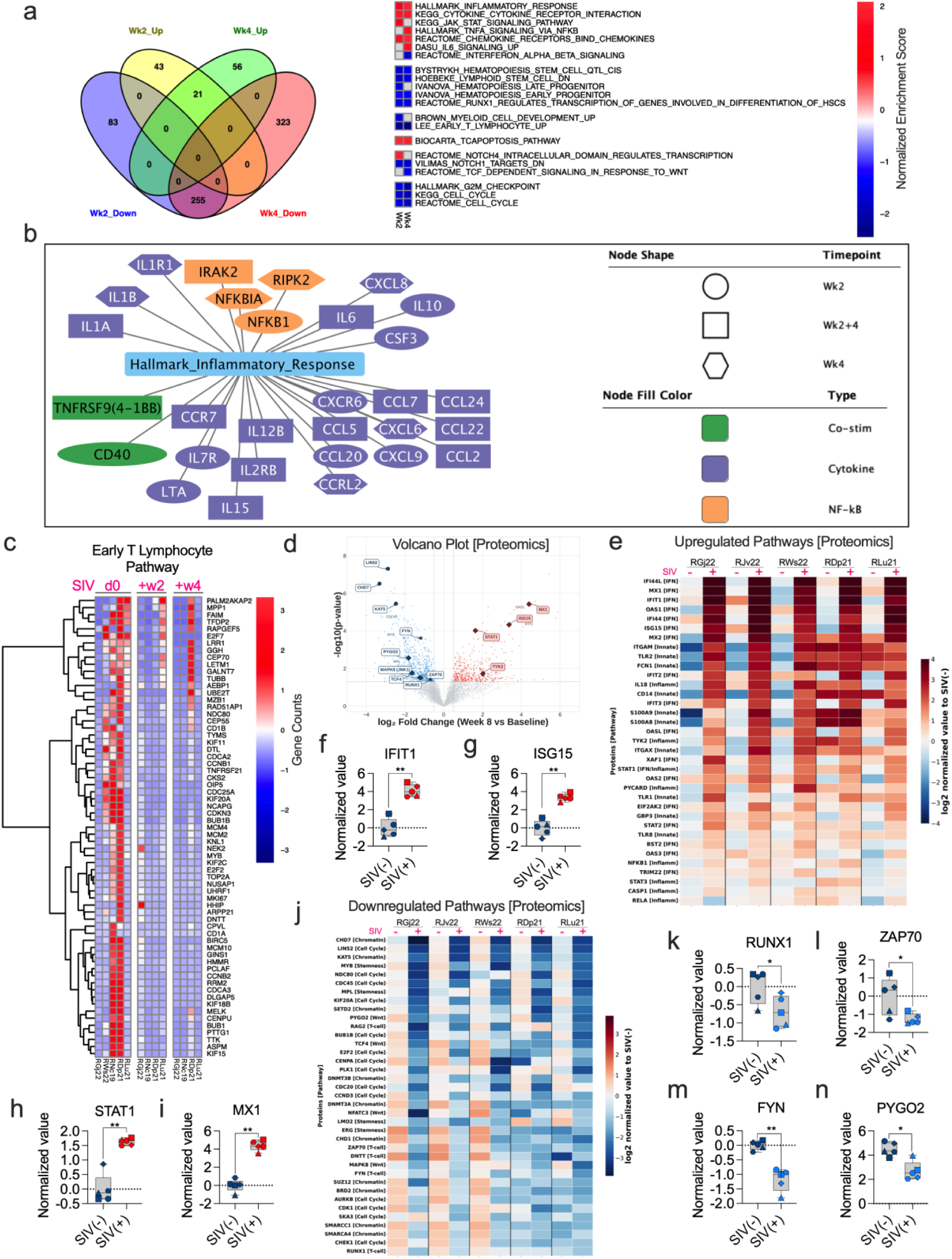
Bone marrow derived HSPCs demonstrated transcriptomics and proteomics signatures associating with increased inflammatory and reduced T cell differentiation pathways (a-n). **a,** Differential gene expression and pathway enrichment analysis performed on total RNA sequencing of sorted CD34⁺ hematopoietic stem and progenitor cells (HSPCs) isolated from bone marrow before infection (SIV⁻) and at 2 and 4 weeks post SIV infection. Venn diagram shows overlapping and distinct transcriptional signatures at week 2 (Wk2) and week 4 (Wk4) relative to baseline. Heatmap displays normalized enrichment scores (NES) from gene set enrichment analysis (GSEA) for selected pathways, revealing increased inflammatory and interferon-associated responses alongside reduced hematopoietic differentiation and cell-cycle programs. **b,** Network visualization of leading-edge genes from the *Hallmark Inflammatory Response* pathway significantly upregulated at Wk2 and/or Wk4. Nodes represent individual genes, with node shape indicating time point and node color denoting functional category, highlighting induction of NF-κB signaling components, cytokines, chemokines, and co-stimulatory molecules. **c,** Row-normalized heatmap showing reduced expression of leading-edge genes from the *Lee_Early_T_Lymphocyte_Up* gene set at Wk2 and Wk4, indicating suppression of early T-cell differentiation programs in bone-marrow HSPCs following SIV infection. **d,** Volcano plot of proteomic data derived from bone-marrow CD34⁺ HSPCs isolated from five rhesus macaques (n = 5), comparing week 8 post SIV infection with baseline. The x axis represents log₂ fold change (Week 8/Baseline), and the y axis represents −log₁₀(P value) from two-sample t-tests. Vertical dashed lines denote ±0.5 log₂ fold-change thresholds, and the horizontal dashed line indicates the significance threshold (P = 0.05). Significantly upregulated proteins (log₂FC > 0.5, P < 0.05; n = 465) are shown in red, significantly downregulated proteins (log₂FC < −0.5, P < 0.05; n = 657) in blue, and nonsignificant proteins in grey. Highlighted downregulated pathways include T-cell lineage regulators (FYN, ZAP70, RUNX1; star symbols), Wnt/β-catenin signaling components (PYGO2, TCF4, MAPK8/JNK1; diamond symbols), and cell-cycle or chromatin regulators (CHD7, LIN52, KAT5; hexagon symbols). Upregulated factors (dark red diamonds) include components of the type I interferon and JAK–STAT pathways (MX1, ISG15, STAT1, TYK2). Additional interferon-stimulated genes (IFIT1, OAS1) and stemness/proliferation-associated proteins (MYB, MPL, CDC45) are annotated. **e,** Heatmap showing log2 fold change compared to baseline (week 0 of SIV infection) expression of 35 upregulated proteins across five rhesus macaques at baseline [SIV(−)] and week 8 post infection [SIV(+)]. Samples are arranged by individual animal (RGJ22, RJV22, RWS22, RDP21, RLU21; left to right), with paired pre- and post-infection samples. Vertical black lines delineate individual animals. Proteins are ordered by decreasing log₂ fold change. Color scale represents row-wise Z-score normalization (−2.5 to +2.5). Functional annotations are indicated in brackets: [IFN], type I interferon response; [Inflamm], inflammasome/IL-6/JAK–STAT signaling; [Innate], innate immune and pattern-recognition pathways. **f–i,** Box plots of representative inflammation associated upregulated proteins: IFIT1 (f), ISG15 (g), STAT1 (h), and MX1 (i). **j,** Heatmap showing log2 fold change compared to baseline (week 0 of SIV infection) of 36 downregulated proteins across the same five animals at baseline and week 8 post SIV infection, arranged and normalized as in **e**. Proteins are ordered by increasing log₂ fold change (most downregulated first) and include chromatin modifiers (CHD7, KAT5, SETD2, DNMT3A/B, SUZ12), cell-cycle regulators (LIN52, CDC45, PLK1, CDK1, CCND3), Wnt/β-catenin pathway components (PYGO2, TCF4, MAPK8), T-cell lineage factors (RAG2, ZAP70, DNTT, FYN, RUNX1), and stemness-associated regulators (MYB, MPL, LMO2, ERG). **k–n,** Box plots of representative T cell associated downregulated proteins: RUNX1 (k), ZAP70 (l), FYN (m), and PYGO2 (n), in bone-marrow CD34+ HSPCs at baseline [SIV(−)] and week 8 post infection [SIV(+)]. Each symbol represents an individual animal. Data are shown as median, minimum, and maximum value. Statistical significance was determined using paired two-tailed tests. *P < 0.05; **P < 0.01.

### Transcriptomic and proteomic profiling revealed inflammatory imprinting of HSPCs antagonizing T cell developmental pathways

To define transcriptional signatures modulated by SIV infection, we performed RNA sequencing on sorted CD34⁺ HSPCs from BM at day 0 (n=5) and weeks 2, 4, and 8 p.i. (n=4). Gene set enrichment analysis (GSEA) revealed ∼17.5% (21 of 120 pathways) of induced pathways at both Wk2 and Wk4 post infection, whereas 255 of 661 (∼38.5%) of reduced pathways were common to Wk2 and Wk4, indicating more conservation of reduced pathways in the bone marrow (Fig. 5a). Importantly, there was no overlap between induced and suppressed pathways at either Wk2 or Wk4. Induced pathways included Hallmark Inflammatory Response, cytokine/chemokine signaling, TNF-α via NF-κB, and JAK/STAT/IL-6 signaling, indicating highly inflammatory state of CD34⁺ HSPCs during acute infection (Fig. 5a). Leading-edge analysis identified critical NF-κB signaling components (NFKB1, NFKBIA, IRAK2) and inflammatory cytokines (IL-1β, IL-6, CXCL8) (Fig. 5b). Concurrently, we observed significant reduction in pathways of hematopoiesis, T lymphocyte development, and NOTCH signaling (Fig. 5c), with decreased expression of genes regulating cellular division (MKI67, CDCA2, CDC25A), transcriptional regulation (E2F7, E2F2), and apoptosis inhibition (FAIM) (Fig. 5c). Reduced expression of FAIM links with the increased expression of the TCR apoptosis pathways was also observed at Wk2 and Wk4 (Fig. 5a). Together, these results suggest rapid inflammatory response driven changes in bone marrow HSPCs as early as week 2 post infection and negatively regulates pathways of hematopoiesis, T cell development, and cell cycling.

Next, mass spectrometry-based proteomic analysis of BM CD3⁻CD34⁺ HSPCs (n=5) at week 8 p.i. compared to baseline identified 465 significantly upregulated and 657 downregulated proteins (P<0.05, |log₂FC|>0.5) (Fig. 5d). Key inflammatory mediators were significantly elevated, including IL-18 (+2.40 log₂FC), PYCARD/ASC (+1.55 log₂FC), TYK2 (+2.01 log₂FC), and STAT1 (+1.63 log₂FC), alongside robust induction of interferon-stimulated genes (IFI44L, +5.42 log₂FC; MX1, +4.39 log₂FC; ISG15, +3.36 log₂FC) (Fig.5e-i).

Notably, proteomic profiling revealed impaired T cell lineage programming, with significant downregulation of RUNX1 (−0.69 log₂FC, P=0.038), FYN (−1.18 log₂FC), ZAP70 (−1.27 log₂FC), and RAG2 (−1.81 log₂FC) (Fig. 5j-m). Wnt/β-catenin signaling components were also suppressed (PYGO2, −1.81 log₂FC; TCF4, −1.63 log₂FC), alongside stemness-associated transcription factors (MYB, −2.45 log₂FC; LMO2, −1.34 log₂FC) (Fig. 5n). These data indicate SIV infection induces inflammatory imprinting characterized by IL-6/JAK-STAT pathway activation coupled with negative regulation of cell cycle and T cell lineage differentiation potential.

### IL-6 exposure enforced a CD38⁻ progenitor phenotype and impaired DN1-to-DN2 (TSP) transition in rhesus- and human-specific models of thymopoiesis

Given our in vivo results demonstrating IL-6 association with decreased T cell lineage commitment, we leveraged our artificial thymic organoid systems to directly investigate IL-6’s role in antagonizing T cell lineage bias(21). RhATO cultures treated with IL-6 (dose ranging from 0.5–50 ng/mL) showed significant dose-dependent decrease (P = 0.002) in the frequency of DN CD34⁺CD7⁺ T-cell committed progenitors (Fig. 6a-b). Analysis of cell number in RhATO exposed to a static 5ng/mL dose of IL-6 significantly reduced (∼3-fold) cell numbers of DN CD34+CD7+ population from ∼2,600 to ∼895 per 5,000 input HSPCs (Supplementary Fig. 7a). IL-6 receptor blockade rescued this effect, increasing DN CD34⁺CD7⁺ TSP frequency to ∼40% compared with ∼18% in IL-6-treated cultures without IL-6R-blockade (vs. ∼53% in untreated controls; Fig. 6c–e). Similarly, frequency of CD4⁺CD3⁻ ISPs showed >2-fold decrease (P = 0.01) under IL-6 treatment (∼16% vs. ∼34% in controls), which was restored with IL-6R blockade (∼29%; Fig. 6f-g).

**Figure 6.**
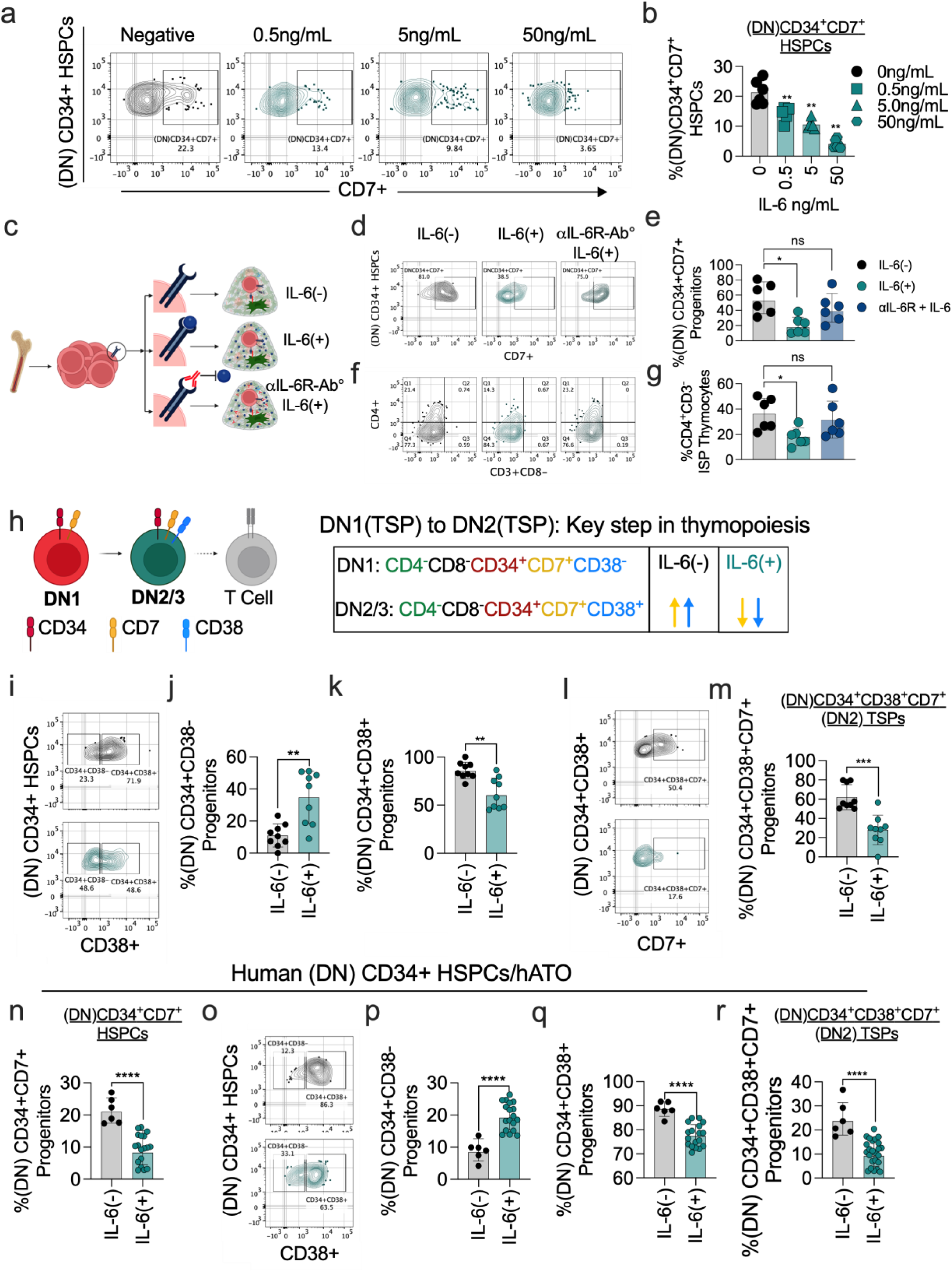
Interleukin-6 impaired T cell committed differentiation of bone marrow derived CD34⁺CD3^−^ HSPCs in a thymus-independent artificial organoid system (a-r). **a**, Representative flow-cytometry plots showing frequencies of DN CD34⁺CD7^+^ T cell committed progenitors in RhATO cultures supplemented without (positive control) or with increasing concentrations of IL-6 (0.5, 5 or 50 ng ml⁻¹), analyzed at week 1 of culture establishment with bone marrow derived CD3^−^CD34^+^ HSPCs from SIV naïve rhesus macaques (RMs) as previously described(21). DN progenitors were identified based on CD4^−^CD8^−^ double negative (DN), CD34 and CD7 expression; numbers indicate frequencies within the gated populations. **b**, Bar graph showing quantification of the frequency of (DN)CD34⁺CD7⁺ TSPs across IL-6 concentrations, demonstrating a dose-dependent reduction in T cell progenitors. **c,** Schematic of the IL-6 receptor blockade experimental design. Bone marrow derived CD3^−^CD34⁺ HSPCs from SIV naive RMs were FACS sorted and cultured RhATO system in the absence of IL-6 (grey), presence of IL-6 (green), and in the presence of IL-6 receptor binding antibody + IL-6 (blue), as indicated. **d**, Representative flow-cytometry plots showing (DN)CD34⁺CD7^+^ T cell progenitors in RhATO cultures at week 1 of culture establishment with bone marrow derived CD3^−^CD34⁺ HSPCs from SIV naive RMs for above-described treatment conditions. **e**, Bar graph showing quantitation of the frequency of (DN)CD34⁺CD7⁺ progenitors across treatment conditions (N=6 RhATOs). **f**, Representative flow-cytometry plots showing T cell lineage committed differentiation of CD3^−^CD34⁺ HSPCs marked by increase in the frequency of CD4^+^CD3^−^ immature single positive thymocytes (ISPs) in RhATO. ISPs (early thymocyte intermediate) were identified based on CD45^+^ CD8^−^ CD3^−^ and CD4^+^ expression; numbers indicate frequencies within the gated populations. **g**, Bar graph showing quantification of the frequency of CD4^+^CD3^−^ ISPs across treatment conditions. **h**, Schematic illustrating key early intermediates along the T cell committed differentiation (CD4^−^CD8^−^ double negative, DN1 (CD38^−^), DN2/3(CD38^+^), immature single-positive thymocyte; ISP) and associated marker expression (CD34, CD7, CD38 and CD4). Arrows indicate relative changes in marker expression under IL-6⁻ and IL-6⁺ treatment conditions. **i**, Representative flow-cytometry plots showing frequencies of CD34^+^CD38^−^ and CD34^+^CD38^+^ fractions within CD4^−^CD8^−^(DN) HSPCs (SIV naïve) cultured in RhATO-system in the absence or presence of IL-6. **J-k**, Bar graphs showing quantitation of the frequencies of (DN)CD34⁺CD38⁻ (j) and (DN)CD34⁺CD38⁺(k) progenitors under IL-6⁻ and IL-6⁺ treatment conditions (N = 9 RhATOs). **l**, Representative flow-cytometry plots showing early T cell lineage committed differentiation to CD34⁺CD38^+^CD7^+^ (DN2) TSPs in RhATO cultures under IL-6⁻ and IL-6⁺ treatment conditions (N = 9 RhATOs). **m**, Bar graph showing quantitation of the frequency of CD34⁺CD38^+^CD7+ (DN2) TSPs under IL-6⁻ and IL-6⁺ treatment conditions. **n-r**, Analysis of the impact of IL-6 exposure on expansion of CD34+CD38+CD7+ (DN2) TSP intermediate in a human-specific nonanimal model of thymopoiesis. **n,** Bar graph showing quantitation of (DN)CD34⁺CD7⁺ T cell progenitors under IL-6⁻ (N = 6 hATOs) and IL-6⁺ (N = 18 hATOs) conditions. **o,** Representative flow-cytometry plots showing CD38 expression within CD34⁺CD7⁺ DN progenitors in hATO culture under IL-6⁻ and IL-6⁺ conditions. **p-q**, Bar graph showing quantitation of the frequencies of CD34⁺CD38⁻ (p) and CD34⁺CD38⁺ (q) progenitors within CD4^−^CD8^−^ (DN) CD34^+^ HSPCs (HIV naïve) under IL-6⁻ and IL-6⁺ conditions. **r**, Bar graph showing quantitation of the frequencies of CD34⁺CD38⁺CD7⁺ (DN2) TSPs under IL-6⁻ and IL-6⁺ conditions. Each point represents an individual organoid. Bar graphs show geomean with geomean standard deviation. Statistical significance was assessed using repeated-measures ANOVA with multiple-comparison correction or Mann Whitney unpaired two-tailed t-tests for unrelated data sets; (*P < 0.05; **P < 0.01; ***P < 0.001; ****P < 0.0001; ns, not significant).

Consistent with in vivo findings that bone marrow CD34+ HSPCs preferentially acquired Lin^−^CD38⁻ phenotype under SIV infection, RhATO cultures treated with 5ng/mL IL-6 induced significant shift toward CD38⁻CD34⁺ HSPCs phenotype within a week of culture, increasing approximately 3-fold from 10% to 30% (Fig. 6h–j), with corresponding decrease in CD38⁺CD34⁺ frequency from 84% to 60% (Fig. 6k) at week 1 post culture. These phenotype shifts were reflected in parallel decrease in total cell number (Supplementary Fig. 7b-c). Notably, frequency of CD34⁺CD38⁺CD7⁺ (DN2) TSP decreased 2-fold (P = 0.0008; Fig. 6l-m) under IL-6 exposure, with ∼4-fold reduction in cell numbers (P = 0.004) from ∼2,600 to ∼700 per 5,000 input HSPCs per RhATO system (Supplementary Fig. 7d).

To assess translatability to humans, we utilized human artificial thymic organoid (hATO) cultures(19). We tested human primary CD3^−^CD34^+^ HSPCs from bone marrow of HIV naïve person to generate ATOs. Similar to results obtained in nonhuman primate model, frequency of DN CD34⁺CD7⁺ T cell committed progenitors decreased ∼3-fold (P = 0.001) from ∼21% to ∼8% under IL-6 treatment (Fig. 6n; Supplementary Fig. 7e–g). IL-6 induced 2-fold increase (P = 0.001) in DN CD38⁻CD34⁺ progenitor frequency from ∼8% to ∼19% (Fig. 6o, p), with corresponding decrease in DN CD38⁺CD34⁺ frequency from 89% to 77% (Fig. 6q; Supplementary Fig. 7h, i). Importantly, frequency of CD34+CD38+CD7+ (DN2) TSPs showed 3-fold decrease (P = 0.0008), with ∼3-fold reduction in cell numbers (P = 0.001) from ∼800 to ∼287 (Fig. 6r; Supplementary Fig. 7i, j). These results demonstrate that IL-6 shifts progenitors into an undifferentiated CD38⁻ state, inhibiting DN1-to-DN2 TSP transition and restricting T cell lineage-biased differentiation.

## Discussion

This study identifies a previously unrecognized pre-thymic mechanism by which inflammation-driven emergency hematopoiesis restricts T cell lineage commitment and associates with viral control early during chronic lentiviral infection before an overt hematologic decline has been reported (44). Using longitudinal analysis of bone marrow hematopoiesis in the SIV–rhesus macaque model of HIV/AIDS, we provide the first in vivo evidence that T cell lineage commitment of bone marrow hematopoietic stem and progenitor cells (HSPCs) is disrupted early during persistent latent infection and remains incompletely restored during chronic stage. These findings reposition bone marrow–derived thymus-seeding progenitors (TSPs) as critical upstream determinants of immune reconstitution and antiviral T cell responses.

We identified a conserved CD4⁻CD8⁻CD34⁺CD38⁻CD7⁺ progenitor population in bone marrow that phenotypically and functionally corresponds to DN1-stage TSPs described in humans(17, 42). The abundance of these progenitors closely tracked thymic T cell–committed progenitors and directly dictated their ex vivo T cell differentiation capacity, establishing TSPs as an obligate pre-thymic checkpoint linking bone marrow hematopoiesis to thymic output. Notably, TSPs declined sharply within two weeks of SIV infection—well before reported thymic involution(6, 45). This temporal dissociation challenges prevailing models that primarily attribute defective T cell reconstitution to thymic damage, instead supporting a paradigm in which early restriction of T cell–biased progenitors initiate durable immune deficits that precede and potentially drive loss of viral control. Direct viral cytopathy is unlikely to account for TSP loss, because: 1) SIV is a CCR5-tropic virus and lacks CXCR4 tropism(46), a prominent chemokine expressed by progenitors, and 2) TSPs are CD4⁻CD8⁻ double-negative cells lacking canonical viral entry receptors(17, 47).

Mechanistically, our data identify interleukin-6 (IL-6) as a principal driver of TSP attrition likely due to its reported role in promoting emergency hematopoiesis, known to favor myeloid over lymphoid lineage biased differentiation of HSPCs (29–31). Consistently, peak IL-6 during acute infection (week 2 post infection) associated with maximal disruption of T cell lineage commitment. In reductionist ex-vivo models of thymopoiesis, IL-6 exposure of HSPCs was sufficient to impair T cell commitment, an effect reversible with IL-6 receptor blockade, directly implicating IL-6 signaling as a modifiable determinant of T cell lineage bias. Longitudinal analyses further revealed that progenitor fate restriction was dynamic rather than static. At two weeks post-infection, peak viremia and inflammation coincided with expansion of total CD34⁺ HSPCs alongside a marked loss of TSPs and reduced T cell differentiation capacity, consistent with emergency hematopoiesis rather than stem cell depletion. Partial resolution of inflammation by week four was associated with modest recovery of lineage-primed progenitors, but HSPCs remained persistently biased toward a lineage-negative state throughout chronic infection, indicating incomplete reversal of early inflammatory imprinting. This pattern aligns with established paradigms in which inflammatory cytokines suppress lymphoid differentiation in favor of myelopoiesis as an adaptive stress response(30, 33, 34). Consistent with this framework, we also observed early IP-10 (CXCL10) induction, a chemokine ligand for CXCR3, implicated in myeloid skewing and progenitor fate redirection(43). Consistently, we observed strong association between plasma IP-10 and decrease in TSPs, which further support the idea that a myeloid biased hematologic response antagonizing T cell lineage commitment of bone marrow progenitors under SIV infection.

Notably, CD38 emerged as a key molecular checkpoint integrating inflammatory signaling with T cell lineage commitment progression. CD38 upregulation marks the transition from DN1 to DN2 TSPs and is required for subsequent thymocyte differentiation(17, 48). We show that IL-6 suppresses CD38 expression on HSPCs, enforcing retention in a primitive, undifferentiated state and creating a developmental bottleneck at the earliest stage of thymopoiesis. In both rhesus and human artificial thymic organoid systems, IL-6 exposure reproducibly increased the frequency and number of CD38⁻ progenitors while collapsing CD38⁺ DN2-stage TSPs, demonstrating cross-species conservation of this mechanism. The concordance between in vivo primate data and human-relevant ex vivo platforms eliminate confounding effects of systemic infection and underscores the robustness of the IL-6–CD38 axis as a regulator of T cell lineage commitment. Further investigation of the molecular axis underlying this inflammation driven CD38 downregulation accompanying T cell lineage antagonism is warranted.

These phenotypic findings were further supported by molecular profiling. Transcriptomic and proteomic analyses revealed inflammatory imprinting of HSPCs characterized by downregulation of core T cell developmental regulators, including RUNX1, ZAP70, and FYN. Although these analyses were performed on bulk populations and limited timepoints, the observed signatures were consistent with impaired lymphoid programming and limited cell cycling. Consistently, earlier work has shown reduced differentiation of CD34+ HSPCs, isolated from chronically SIV infected nonhuman primate, to CD34+CD7+ T cell progenitors in ex-vivo cultures and reported arrested G0/G1 cell cycle phase of cells (49). While this study did not investigate CD38 phenotype, it is likely that progenitors were enforced in CD38^−^CD34^+^ state consistent with prior studies reporting CD38⁻CD34⁺ HSPCs preferentially adapting G0/G1 phase(50). Attempts to define immune potential of HSPCs under HIV/AIDS conflate adaptive emergency responses with intrinsic stem cell dysfunction by failing to account for the physiological tendency of HSPCs to restrict lymphoid differentiation under inflammatory stress. Our data instead indicate that impaired T cell regeneration arises from inflammation-driven enforcement of an undifferentiated progenitor state rather than irreversible hematopoietic damage. Transcriptional and proteomic profiling substantiated this model revealing inflammatory imprinting characterized upregulation of inflammatory pathways and a parallel downregulation of core T cell developmental regulators including decreased cell cycling. The molecular signatures were consistent with phenotypic and functional data. Future single-cell epigenetic and transcriptomic profiling of bone marrow and thymic progenitors will define lineage-specific trajectories and determine how inflammatory imprinting differentially constrains distinct progenitor subsets underlying mature T cell reconstitution.

Importantly, early pre-thymic restriction had durable consequences: TSP abundance at two weeks strongly predicted ex vivo T cell output from HSPCs isolated at later timepoints and directly associated with loss of viral control, independent of peripheral T cells. The magnitude of immune failure is thus established remarkably early and persists beyond the initial inflammatory insult under chronic antigen persistence. Given antiretroviral therapy (ART) does not completely resolve inflammation, it’s imperative to understand IL-6 driven inhibition of T cell lineage commitment of bone marrow HSPCs under ART both in NHP model (critical for pre-clinical HIV cure studies) and in people living with HIV. The clinical implications of our findings are substantial. Early TSP depletion predicted impaired T cell regeneration and higher viral load across all timepoints, providing mechanistic explanation for persistent immune dysfunction observed in individuals despite suppressive antiretroviral therapy (ART)(40, 51–54). Chronic low-grade inflammation with elevated IL-6 persists in many ART-treated individuals; our findings suggest this residual inflammatory milieu may continue impairing T cell lineage bias, limiting immune reconstitution independently of viral replication. This framework aligns with clinical phenotypes in which immunological non-responders exhibit elevated IL-6, reduced thymic output, and poor CD4⁺ T cell recovery, whereas elite controllers maintain low inflammation and preserved immune regeneration. Intriguingly, in a nonpathogenic model of lentiviral infection, efficient CD4 T cell reconstitution is associated with low inflammatory IL-6 induction despite persistent SIV infection(55, 56). These clinical and pre-clinical evidence further substantiate our findings.

Beyond immune reconstitution, early restriction of T cell regenerative capacity likely exacerbates viral reservoir establishment by limiting timely antigen-specific containment during acute infection. Collectively, these findings argue that durable immune restoration requires not only viral suppression but targeted modulation of inflammation to restore T cell–biased hematopoiesis. Therapeutic strategies targeting IL-6/IL-6R or downstream JAK–STAT signaling warrant evaluation for preserving bone marrow T cell potential and synergizing with cure interventions. By identifying pre-thymic progenitor fate as a central, modifiable determinant of viral control, this study reveals a previously unrecognized axis governing antiviral immunity that may extend to other chronic inflammatory conditions where sustained inflammation restricts T cell lineage commitment and limits immune surveillance.

## Materials and Methods

### Rhesus macaque study design

Six Indian-origin, specific-pathogen-free Rhesus macaques (RMs) also known as *Macaca mulatta* were recruited from the Emory National Primate Research Center (ENPRC). All RMs were single-housed in an animal BSL-2 facility at ENPRC and provided with both standard primate feed (Jumbo Monkey Diet 5037; Purina Mills, St. Louis, MO), fresh fruit, and water along with daily enrichment. RMs were between 30 and 90 months old at the time of infection. All RMs were *Mamu*-B*07^−^, and *Mamu*-B*17^−^. Three RMs were *Mamu*-A*01^+^ (Supplementary Table 1). RMs were infected with 300 TCID_50_ SIVmac_239M_ (kindly provided by Dr. Brandon Keele, Frederick National Laboratory for Cancer Research, NCI/NIH) via intravenous route. Bone marrow hematopoietic stem and progenitor cells (HSPCs), markers of inflammation in plasma, and peripheral T cells in blood were longitudinally followed until week 16 post infection. Samples were longitudinally collected at various timepoints (Fig. 1s). sample collections were performed under anesthesia with ketamine (5 to 10 mg/kg) or telazol (3 to 5 mg/kg) by trained research and veterinary staff. Additional samples from fifteen SIV naïve and six SIV infected RMs were procured through Emory National Primate Research Center Biological Materials Procurement Program following approved protocols.

### Bioethical approvals

Bone marrow aspirates were obtained from rhesus macaques through the Emory National Primate Research Center Biological Materials Procurement Program, in accordance with the Animal Welfare Act and the National Institutes of Health (NIH; Bethesda, MD) Guide for the Care and Use of Laboratory Animals. All procedures involving nonhuman primates were approved by the Emory University Institutional Animal Care and Use Committee (IACUC) under protocol IPROTO202400000081 (approved August 9, 2024). Bone marrow aspirates were collected from young adult rhesus macaques. Human bone marrow derived CD34^+^ hematopoietic stem and progenitor cells (Cat #BM34C-5) were purchased from Charles River Laboratory Inc, Wilmington, MA, USA.

### Processing of rhesus macaque thymus tissue

Intact thymic tissue from eight Rhesus macaques was processed to generate single cell thymocyte suspensions. Briefly, thymus tissue was dissected into small fragments using a sterile scalpel in a 10-cm^2^ culture dish and further minced with sterile scissors. Thymocytes were released by gently pressing the tissue fragments against the dish using a sterile syringe plunger, followed by mechanical dissociation through repeated passage through an 18-gauge needle to disrupt remaining aggregates. The resulting cell suspension was filtered through a 100-µm cell strainer and washed with PBS by centrifugation at 300 × g for 5 min. The supernatant was discarded, and the cell pellet was resuspended in 3 mL ACK red blood cell lysis buffer (Quality Biological, Cat. no. 118-156-101) and incubated for 3–4 min at room temperature. Lysis was quenched by dilution with FBS-supplemented PBS to a final volume of 25 mL. Cells were washed twice with FBS-supplemented PBS, resuspended in an appropriate buffer, counted, and either used immediately for downstream applications or stored as indicated. Unless otherwise specified, all steps were performed on ice.

### Processing of rhesus macaque peripheral blood mononuclear cells

Blood samples were collected in Vacutainer CPT™ mononuclear cell preparation tubes and centrifuged at 1,500 × g for 30 min according to the manufacturer’s instructions. Following centrifugation, plasma was removed, and the mononuclear cell layer was transferred to a sterile 50-ml conical tube. Cells were washed with PBS by centrifugation at 300 × g for 5 min. The supernatant was discarded, and the cell pellet was resuspended in 3 ml ACK red blood cell lysis buffer and incubated for 3–4 min at room temperature. Lysis was quenched by dilution with FBS-supplemented PBS to a final volume of 25 ml, followed by centrifugation at 300 × g for 5 min. Cells were washed once more with FBS-supplemented PBS, resuspended in an appropriate buffer, counted, and either used immediately for downstream applications or stored as indicated.

### Processing of rhesus macaque bone marrow derived biospecimen

Bone marrow aspirates were obtained from young adult rhesus macaques, including SIV-negative animals and animals infected with SIV and sampled at defined time points following infection, either through the Emory National Primate Research Center Biological Materials Procurement Program or from ongoing IACUC-approved studies. Human mobilized peripheral blood was obtained as discarded material following cell-therapy procedures at Emory Clinic. Bone marrow aspirates (4–5 ml) were diluted 1:1 with PBS supplemented with 2% FBS and passed through a 100-µm cell strainer. Filtered bone marrow aspirates were layered directly over 15 ml Lymphoprep (STEMCELL Technologies, Cat. no. 07851) in SepMate™ conical tubes (STEMCELL Technologies, Cat. no. 85450). Bone marrow scrap (BMS) was resuspended in 25 ml PBS supplemented with 2% FBS, filtered through a 100-µm cell strainer, further diluted to 50 ml, and split into two 25-ml fractions, each layered over 15 ml Lymphoprep in standard conical tubes (Corning, Cat. no. 430921). Samples were centrifuged at 1,500 × g for 20 min at room temperature. The mononuclear cell layer was collected into a separate 50-ml tube and washed with FBS-supplemented PBS by centrifugation at 300 × g for 5 min. Red blood cells were lysed by resuspending the cell pellet in 4 ml ACK lysis buffer and incubating for 3–4 min at room temperature. Lysis was quenched by dilution with FBS-supplemented PBS to a final volume of 30–50 ml, depending on the initial BMS content, followed by centrifugation at 300 × g for 5 min. Cells were washed twice with FBS-supplemented PBS, resuspended in an appropriate buffer, counted, and either used immediately for downstream applications or stored as indicated. Unless otherwise specified, all buffers and samples were maintained on ice throughout processing.

### Enrichment and isolation of CD3^−^CD34^+^ HSPCs

Rhesus macaque CD34⁺ hematopoietic stem and progenitor cells (HSPCs) were enriched using the Dynabeads™ CD34 Isolation Kit (Invitrogen, Cat. no. 11301D) according to the manufacturer’s instructions. Briefly, 4 × 10⁷ bone-marrow mononuclear cells were incubated with 100 µl Dynabeads on ice for 30 min. Cells were then brought to a final volume of 2 ml using MACS buffer (STEMCELL Technologies, Cat. no. 20144) and placed on a magnetic stand for 2 min to separate bead-bound cells from unbound cells, which were discarded. This magnetic separation step was repeated once, after which bead-bound cells were resuspended in 100 µl buffer. Dynabeads were detached by adding 100 µl DETACHaBEAD reagent and incubating for 45 min. To enhance bead detachment, 3 ml MACS buffer was added, and the suspension was vortexed briefly. The tube was placed on a magnetic stand for 2 min, and the supernatant containing detached CD34⁺ cells was transferred to a new tube. Beads were washed once more with MACS buffer, subjected to magnetic separation, and the supernatant was pooled to maximize cell recovery. Enriched CD34⁺ cells were washed with 10 ml MACS buffer to remove residual DETACHaBEAD and either used immediately or stored as indicated. For fluorescence-activated cell sorting (FACS), enriched cells were stained with a viability dye and antibodies against CD34, CD3, CD4, CD8, and CD7 markers. For characterization of (DN)CD34^+^CD7^+^ TSPs, CD4^−^CD8^−^CD3^−^CD34^+^CD7^+^ progenitors were sorted and RhATOs were established using 5000 input progenitor cells. T cell lineage biased differentiation was analyzed at week 4 of culture. For routine investigation of T cell lineage committed differentiation of bone marrow progenitors upon SIV infection, live CD3⁻CD34⁺ cells were FACS sorted from bone marrow mononuclear cells and RhATO cultures were established using 5000 input cells. Analysis was performed at week 1 or week 4 of culture depending upon the thymic intermediate analyzed. <u>Isolation of human CD3-CD34+ HSPC:</u> Total CD34+ HSPCs (Cat#BM34C-5) purchased from Charles River Laboratory Inc, Wilmington, MA, USA, were thawed and directly stained with live/dead stain, anti-CD3, and anti-CD34 antibodies. CD3+ cells were depleted by FACS sorting and CD34+CD3-cells were used to established ATOs immediately after the sort.

### Establishment of rhesus- and human-specific artificial thymic organoids (RhATO/hATO)

Rhesus artificial thymic organoids (RhATO) cultures were established as previously described(21). Similarly, human artificial thymic organoids (ATO) were established as previously described (19). Briefly, all ATO (RhATO/hATO) culture medium consisted of RPMI 1640 supplemented with 1X B27 serum-free supplement (Thermo Fisher Scientific, Cat. no. 17504001), 1% GlutaMAX (Thermo Fisher Scientific, Cat. no. 35050061), 1% penicillin–streptomycin (Thermo Fisher Scientific, Cat. no. P4458), 5 ng ml⁻¹ stem cell factor (PeproTech, Cat. no. 300-07-100UG), 50 µM L-ascorbic acid (MilliporeSigma, Cat. no. A8960), 7.5 ng ml⁻¹ (5 ng ml⁻¹ for hATO) recombinant IL-7 (PeproTech, Cat. no. 200-07-2UG), and 7.5 ng ml⁻¹ (5 ng ml⁻¹ for hATO) FLT3 ligand (PeproTech, Cat. no. 300-19-2UG). Six-well plates were prepared with 1 ml of ATO culture medium per well, and 0.4-µm Millicell™ transwell inserts (Millipore, Cat. no. PICM0RG50) were placed into each well. MS5-RhDLL1 (or MS5-hDLL1 for hATO) cell lines were trypsinized, filtered through a 30-µm cell strainer to remove aggregates, resuspended in culture medium, and counted. Sorted CD3⁻CD34⁺ HSPCs (3,000–7,500 cells) were aggregated with Rh-DLL1 or hDL1 cells at a 1:20 ratio by gentle pipetting. Cell aggregates were centrifuged at 300 × g for 3 min, the supernatant was carefully aspirated, and pellets were resuspended in 5 µl culture medium per ATO. To establish ATOs, 5 µl of each cell aggregate was carefully deposited onto the transwell insert. Cultures were maintained in a humidified incubator at 37 °C with 5% CO₂. Medium was replaced every 3–4 d, and ATOs were cultured for 1–4 weeks, depending on experimental requirements and desired thymocyte yield. For analysis of early intermediates (CD34^+^CD7^+^ TSPs, CD34^+^CD38^+^CD7^+^ (DN2) TSPs, and CD4^+^CD3^−^ immature single positive thymocytes (ISPs) RhATOs were processed and analyzed at week 1 post culture establishment. For analysis of CD3^+^ T cell development, ATOs were analyzed at week 4 post culture establishment. <u>IL-6 treatment experiments</u>: To analyze dose dependent impact of IL-6, ATOs were maintained under increasing culture concentrations (0.5, 5, 50 ng/mL). For other experiments, ATOs were maintained in culture media supplemented with static 5ng/mL of IL-6. <u>IL-6 receptor blockade experiments:</u> Sorted CD3^−^CD34^+^ HSPCs were first incubated with anti-IL-6R antibody for 20 minutes on ice followed by their culture in media supplemented with IL-6 (5ng/mL) and anti-IL-R antibody. The T cell lineage committed differentiation was assessed at week 1 of culture.

### Phenotypic Analysis by Flow Cytometry

To analyze early intermediates of T cell lineage committed differentiation, ATOs were processed at week 1 of culture. To analyze complete T cell differentiation, ATO cultures were analyzed at week 4 of culture. Individual ATOs were processed by adding 1 ml FACS buffer directly onto the transwell insert and gently disrupting the organoid using a pipette tip. The disrupted aggregates were pipetted repeatedly to facilitate release of lymphocytes into suspension. Cell suspensions containing lymphocytes and residual stromal cells were filtered through a 100-µm cell strainer, and surface immunostaining was performed. Cells were stained in FACS buffer with a viability dye and antibodies against CD45, CD34, CD38, CD3, CD8, CD4, CD7 to delineate T-cell developmental stages. To analyze differentiated and primitive CD34+ hematopoietic stem and progenitor cells (HSPCs), bone marrow derived mononuclear cells were stained with viability dye, and antibodies against lineage markers (CD34, CD2, CD3, CD7, CD10, CD11b, CD14, CD19, CD20, CD33, CD56, CD235a) in addition to CD38, and Ki67. For analysis of CXCR3 expression, bone marrow nuclear cells were stained with viability dye, and antibodies against CD34, CD38, CD45, and CXCR3. For analyzing lymphoid primed multipotent progenitors (LMPPs), bone marrow mononuclear cells were stained with viability dye, and antibodies against CD34, CD38, CD45RA, CD90, in addition to lineage markers. For analysis of T cell committed progenitors, bone marrow mononuclear cells and thymocytes were stained with viability dye, antibodies against CD45, CD34, CD38, CD3, CD4, CD8, CD7, TCF1/7 (used only for thymocytes) markers. For intracellular markers such as Ki67, and TCF-1/7, cells were fixed and permeabilized with the eBioscience FoxP3/Transcription Factor staining buffer set following the manufacturer’s instructions and stained with antibodies specific to Tcf1/7, and Ki67. Data were either acquired on a BD FACS Symphony or the Cytec spectral flow cytometer and analyzed using FlowJo software (v10).

### Antibodies

Antibodies used in this study included CD3 (clone SP34-2, BD Biosciences), CD45 (clone D058-1283 (RUO), BD Biosciences), CD8 (clone SK1, BD Biosciences), CD4 (clone L200, BD Biosciences), CD34 (clone 561, BioLegend), CD38 (clone AT1), CD7 (clone M-T70, BD Biosciences), CD90 (5E10, BD Bioscience), CD45RA (clone 5H9, BD Bioscience), TCF1/7 (clone C63D9, Cell Signaling Technology), Ki-67 (clone B56, BD Bioscience), CXCR3 (clone G025H7, BioLegend), CD2 (clone RPA-2, BD Bioscience), CD11b (clone ICRF44, BD Biosciences), CD14 (clone M5E2, BioLegend), CD19 (clone HIB19, BD Biosciences), CD20 (clone 2H7, BD Biosciences), CD10 (clone HI10a, BioLegend), CD33 (clone P67.6, BioLegend), CD235a (clone HI264, BioLegend), CD56 (clone NCAM16.2, BD Biosciences).

### Determination of Plasma Viral RNA

Plasma viral load was quantified as previously described(57). Viral RNA was extracted from 200 µl rhesus macaque plasma using the QIAsymphony DSP Virus/Pathogen Mini Kit (QIAGEN) according to the manufacturer’s instructions, with an elution volume of 60 µl. SIV RNA was quantified by one-step quantitative reverse transcription PCR (qRT–PCR) using TaqMan™ Fast Virus 1-Step Master Mix (Applied Biosystems) and SIV *gag*-specific primers and probe (forward: 5′-GCAGAGGAGGAAATTACCCAGTAC-3′, Fisher Scientific; reverse: 5′- CAATTTTACCCAGGCATTTAATGTT-3′, Fisher Scientific; probe: 5′ 6FAM-TGTCCACCTGCCATTAAGCCCGA-TAMRA-3′). Absolute quantification was performed using a standard curve generated from fivefold serial dilutions of SIV *gag* plasmid RNA ranging from 2.11 × 10⁸ to 2.69 × 10³ copies ml⁻¹, run in duplicate on each plate. qRT–PCR reactions were performed on an Applied Biosystems 7500 or QuantStudio™ 3 Real-Time PCR System under the following cycling conditions: 50 °C for 15 min, 95 °C for 2 min, followed by 40 cycles of 95 °C for 15 s and 60 °C for 1 min. Plasma viral RNA concentrations (copies ml⁻¹) were calculated from the standard curve (R² ≥ 0.99) and adjusted for sample dilution where applicable. The assay limit of detection was 60 copies ml⁻¹.

### Bulk RNA Sequencing of CD34^+^ HSPCs

To match the progenitors used in ATO cultures, approximately, 25000 CD3^−^CD34^+^ HSPCs from bone marrow aspirates were FACS sorted at weeks 0, 2, and 4 post SIV infection. Sorted cells were lysed and RNA stored in RNAlater solution for future processing. Once all samples were collected, RNA was extracted using RNeasy spin column purification kits (Qiagen) and quantified by nanodrop. 100-1000 ng of total RNA per sample was then converted into cDNA libraries using Stranded Total RNA Prep with Ribo-Zero Plus (Illumina). Sequencing was performed on the Illumina NovaSeq 6000 systems. Raw sequencing fastq files were then aligned, annotated and QC’d. Differential gene expression and GSEA were performed using the package ‘edgeR’. Heatmaps were generated using ‘pheatmap’ and networks of leading-edge genes visualized using Cytoscape.

### Proteomics of CD34⁺ HSPCs

To match the progenitors used in ATO cultures, approximately, 25000 CD3^−^CD34^+^ HSPCs from bone marrow aspirates were FACS sorted from 5 RMs at weeks 8 post SIV infection CD3⁻CD34⁺ hematopoietic stem and progenitor cells were lysed in buffer containing 50 mM Tris-HCl (pH 8.5), 10 mM TCEP, 40 mM CAA, and 0.2% DDM, boiled at 95 °C for 10 min, and sonicated. Proteins were digested overnight at 37 °C using 500 ng Trypsin/Lys-C Mix (Promega). Digests were acidified with trifluoroacetic acid, and approximately one-sixth of each sample was loaded onto preconditioned EvoTips (EvoSep). Peptides were analyzed on an EvoSep One system coupled to a timsTOF Pro 2 mass spectrometer (Bruker Daltonics) using a 15-cm Aurora Elite CSI column with the 40 SPD Whisper Zoom gradient. Data were acquired in positive ion mode using DIA-PASEF(58), with acquisition parameters optimized by py_diAID(59), employing four ion mobility windows and twelve m/z isolation windows across an m/z range of 100–1700 and ion mobility–dependent collision energies. Protein identification and quantification were performed in Spectronaut (v20) in library-free mode against the *Macaca mulatta* UniProt database (UP000006718)(60). Carbamidomethylation of cysteine was specified as a fixed modification, while methionine oxidation and protein N-terminal acetylation were included as variable modifications. Proteins quantified in at least three replicates in one group were retained; intensities were median-normalized and log₂-transformed. Missing values were imputed by Bayesian PCA imputation(61) and downshift sampling, for proteins missing at random or not at random, respectively(62). Statistical comparisons were performed using two-tailed Student’s *t*-tests, with *P* < 0.05 considered statistically significant.

### Mesoscale discovery multiplex cytokine assay

Multiplex assay was performed as previously described(63). Briefly, A panel of cytokines and chemokines was quantified in plasma using a multiplexed electrochemiluminescence-based assay platform (Meso Scale Discovery, MSD). Proinflammatory mediators were measured using the V-PLEX Proinflammatory Panel 1 (NHP), cytokines using the V-PLEX Cytokine Panel 1 (NHP), and chemokines using the V-PLEX Chemokine Panel 1 (Human and NHP, Gen B). These panels collectively included IL-1β, IL-2, IL-5, IL-6, IL-7, IL-10, IL-12/IL-23p40, IL-15, IL-16, IL-17A, IFNγ, TNFα, GM-CSF, VEGF, CCL2 (MCP-1), CCL4 (MIP-1β), CCL11 (Eotaxin), CCL17 (TARC), CCL22 (MDC), CCL26 (Eotaxin-3), CXCL9, CXCL10, and CXCL11, among others, as specified by the manufacturer. Assays were performed according to the manufacturer’s instructions and the Emory Multiplexed Immunoassay Core (EMIC) standard operating procedures. Plasma samples were diluted as recommended for each panel, and cytokine concentrations were determined from standard curves generated on each plate using manufacturer-supplied calibrators. Plates were read on a MESO QuickPlex SQ 120 instrument, and analyte concentrations were calculated using MSD Discovery Workbench software. All values were corrected for sample dilution and are reported as pg ml⁻¹.

### Statistical analysis

Depending on the data distribution, the difference between any two groups at a given time point was measured using a two-tailed nonparametric Mann–Whitney rank sum test or an unpaired parametric t-test. Comparisons between different time points within a group used a paired parametric t-test. A p-value of less than 0.05 was considered significant. The correlation analysis was performed using the Pearson test. GraphPad Prism version 10 was used for data analysis and statistics

## Author Contributions

S.A.R. conceived and designed the research, brought funding, conducted the research, analyzed data, interpreted results, and wrote the manuscript. S.A., S.N., C.M.B., D.E.G., and J.A.T. conducted the research and analyzed the data. F.J.T.S., reviewed and interpreted the data, B.F.K., provided SIVmac239M, V.V., M.P., and G.S. provided biospecimen from SIV infected rhesus macaques, R.P.J., M.S.A.H., and A.F.C. helped with data interpretations. S.A.R. supervised the research. All authors reviewed and approved the manuscript.

## Institutional Review Board Statement

This study was conducted in accordance with the Animal Welfare Act and the National Institutes of Health (NIH) (Bethesda, MD, USA) Guide for the Care and Use of Laboratory Animals, using protocols approved by the Emory University Institutional Animal Care and Use Committee. Biospecimens from the Rhesus macaques were obtained from the study approved under IACUC number IPROTO202400000081 on 9 August 2024.

## Data Availability Statement

The raw data were generated at the Emory University and will be available upon request from the corresponding author. The mass spectrometry proteomics data files have been deposited to the ProteomeXchange Consortium via the PRIDE partner repository under project accession: PXD073638(64). The bulk RNASeq raw sequencing reads will be deposited into GEO prior to publication and that all code and processed data will be available on Github.

## Acknowledgments

We thank Stephanie Ehnert, Stacey Weissman, and other members of the research tech services team at the Emory National Primate Research Center (ENPRC). We thank Jennifer S. Wood, Rachelle Lauren Stammen, Sherrie Jean, and all current and former veterinarian scientists at the ENPRC. We thank L. Shan, Magdalen Chouinard and Emory Center for AIDS Research (CFAR) virology core for viral RNA measurement and analysis. We thank Jianjun Chang and Emory Multiplexed Immunoassay Core (EMIC) for multiplex cytokine and chemokine measurement and analysis. We thank Dr. Max D. Cooper, Emory University, Atlanta, GA, for reviewing the results and helpful advice. We thank Dr. Daniel Kalman, Emory University, Atlanta, GA, for constructive critiques of the data. We thank Prof. Rafi Ahmed, Emory University, Atlanta, GA, for constructive critiques of the data. We thank Kiran Gill, Ankur Saini, Robert E. Karaffa II, Kametha Fife, Sommer Durham, and members of the flow cytometry cores at the ENPRC and Emory Vaccine Center, Emory University. We thank Kay Lee Summerville and members of the ENPRC Biological Material Procurement program. The content is solely the responsibility of the authors and does not necessarily reflect the official views of the National Institute of Health. This project has been funded in part with federal funds from the National Cancer Institute, National Institutes of Health, under Contract No. 75N91019D00024. The content of this publication does not necessarily reflect the views or policies of the Department of Health and Human Services, nor does mention of trade names, commercial products, or organizations imply endorsement by the U.S. Government.

## Supplementary Figures

**Supplementary Figure 1.**
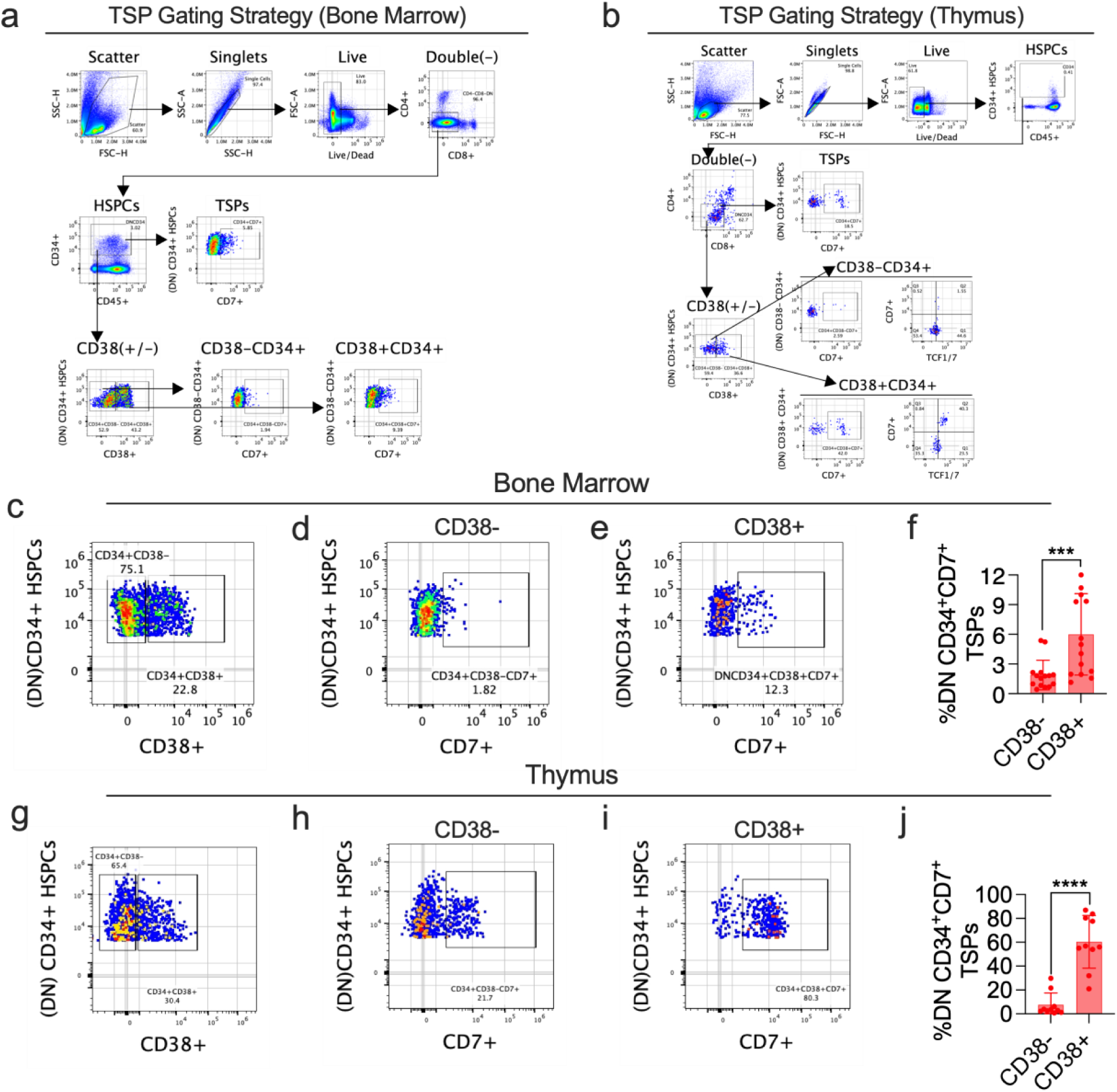
Gating strategy for identification and characterization of thymus-seeding progenitors in bone marrow and thymus (a-j). **a-b,** Flow-cytometric gating strategy used to identify thymus-seeding progenitors (TSPs) within double-negative (DN; CD4⁻CD8⁻) CD34⁺ hematopoietic stem and progenitor cells (HSPCs) in bone marrow (a) and thymus (b). Sequential gating includes scatter properties, singlets, live cells, exclusion of lineage-positive cells (CD4, CD8), and identification of CD34⁺ HSPCs, followed by delineation of CD7⁺ TSPs. DN CD34⁺ cells were further subdivided based on CD38 expression into CD38⁻CD34⁺ and CD38⁺CD34⁺ subsets. In the thymus, CD7⁺ DN CD34⁺ cells were additionally stratified by T-cell factor-1 (TCF-1) expression. **c–e,** Representative flow-cytometric plots from bone marrow showing DN CD34⁺ HSPCs gated by CD38 expression (c), and the distribution of CD7⁺ cells within CD38⁻ (d) and CD38⁺ (e) DN CD34⁺ compartments. **f,** Quantification of the frequency of CD4^−^CD8^−^ double negative (DN) CD34⁺CD7⁺ TSPs within CD38⁻ and CD38⁺ DN CD34⁺ HSPCs in bone marrow. **g–i,** Representative flow-cytometric plots from the thymus showing DN CD34⁺ HSPCs gated by CD38 expression (g), and the distribution of CD7⁺ cells within CD38⁻ (h) and CD38⁺ (i) DN CD34⁺ compartments. **j,** Quantification of the frequency of DN CD34⁺CD7⁺ TSPs within CD38⁻ and CD38⁺ DN CD34⁺ HSPCs in the thymus. Each symbol represents an individual biological sample. Bars indicate geomean ± geomean standard deviation. Statistical significance was assessed using Mann Whitney unpaired two-tailed t-tests. ***P < 0.001; ****P < 0.0001.

**Supplementary Figure 2.**
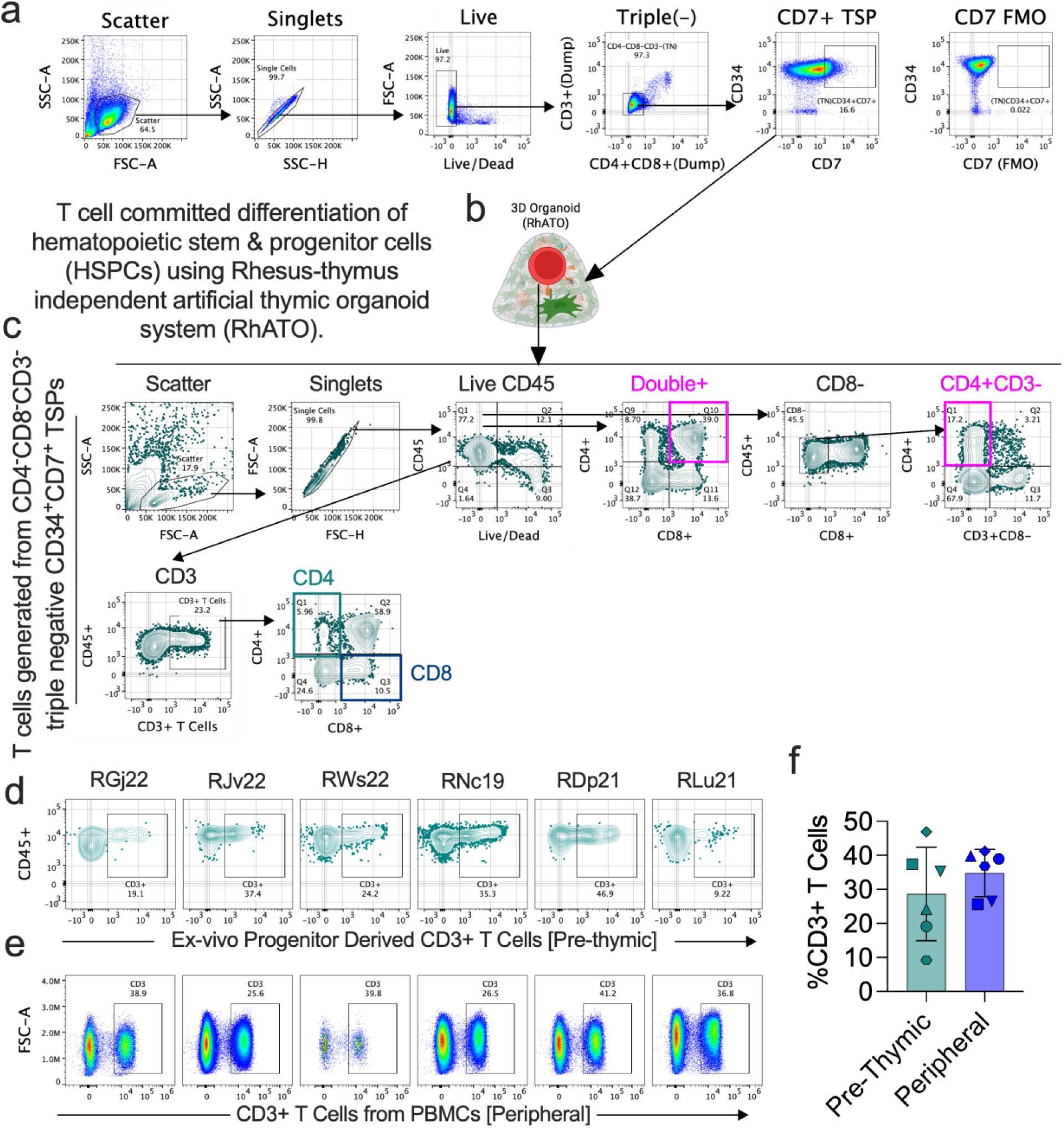
Gating strategy for T cell committed progenitors, their differentiation into T cells in thymus-independent artificial thymic organoid (RhATO) system (pre-thymic T cell potential), and comparison with frequency of peripheral T cells (a-f). **a,** Flow-cytometric gating strategy used to identify CD7⁺ thymus-seeding progenitors (TSPs) from enriched hematopoietic stem and progenitor cells (HSPCs) from bone marrow of SIV naïve Rhesus Macaques (RMs). Sequential gating included scatter properties, singlets, live cells, and exclusion of lineage-positive cells (CD4⁺CD8⁺; dump channel), followed by identification of CD34⁺CD7⁺ triple-negative (CD4⁻CD8⁻CD3⁻) TSPs. Fluorescence-minus-one (FMO) control for CD7 is shown to define gating boundaries. **b,** Schematic illustration of the RM-specific nonanimal model of thymopoiesis known as RhATO-system(21) used to assess T-cell committed differentiation potential of HSPCs isolated from bone marrow of rhesus macaques. **c,** Representative flow-cytometric gating strategy for analysis of thymocyte differentiation following RhATO culture of CD34⁺CD7⁺ TSPs. Cells were gated on singlets, live CD45⁺ cells, followed by identification of double-positive (CD4⁺CD8⁺), CD8⁻ subsets, and CD4⁺CD3⁻ immature thymocytes, as well as mature CD3⁺ T cells, including CD4⁺ and CD8⁺ single positive T cell subsets. **d,** Representative flow-cytometric plots showing CD3⁺ T cells generated in RhATO cultures seeded with HSPCs from bone marrow of SIV-infected RMs at week 16 post-infection, displayed for individual animals (RGj22, RJv22, RWs22, RNc19). **e,** Representative flow-cytometric plots showing CD3⁺ T cells from peripheral blood mononuclear cells of SIV-infected RMs at week 16 post-infection. f, Quantitation of the frequencies of CD3+ T cells generated directly from bone marrow derived HSPCs in RhATO-system (pre-thymic) or PBMCs (peripheral) from same six RMs (N=6). Bar graph showing geomean and individual values.

**Supplementary Figure 3.**
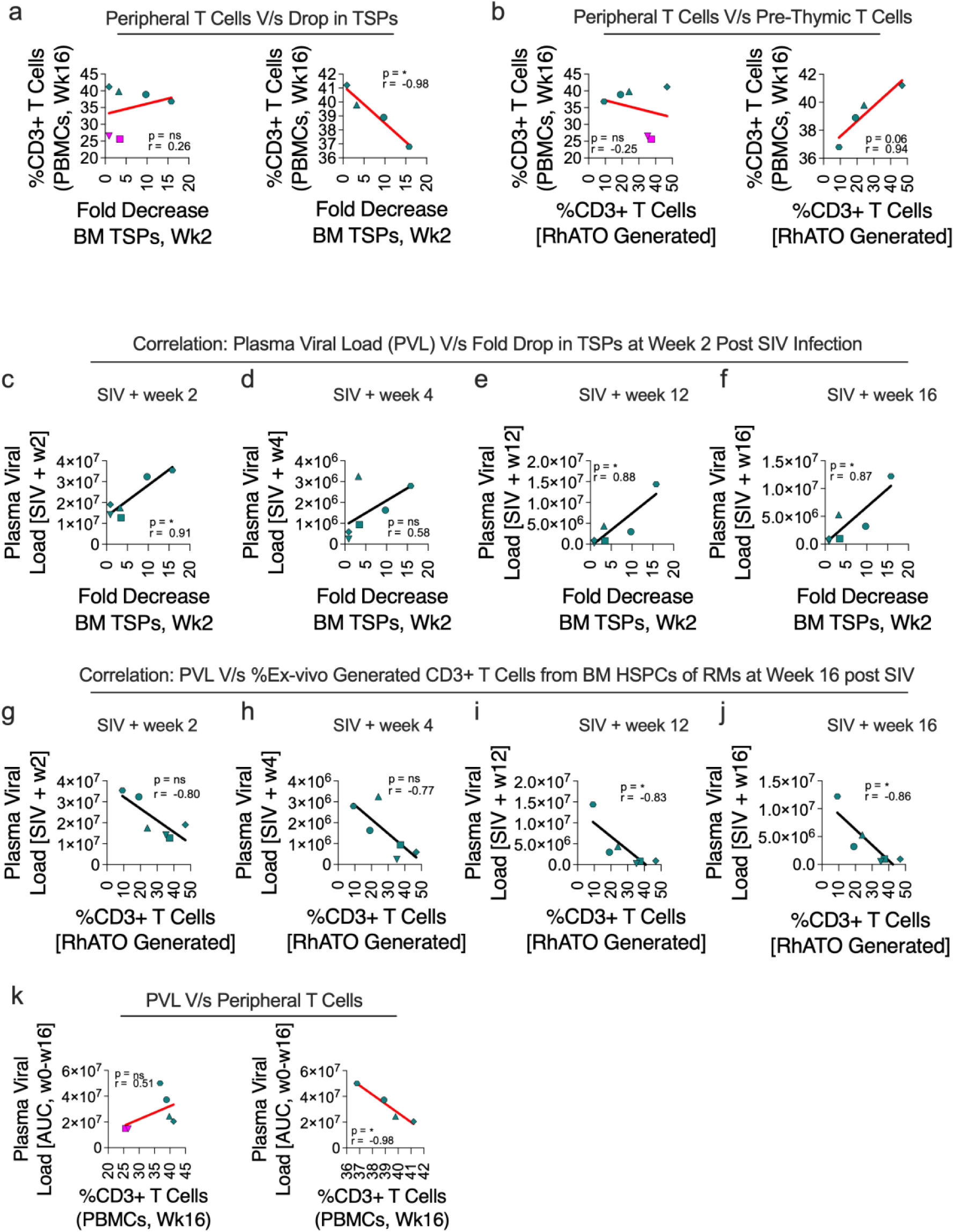
Association of the pre-thymic T cell potential of bone marrow progenitors and peripheral T cells with viral control during SIV infection (a-k). **a,** Correlation between frequency of peripheral T cells at week 16 post SIV infection and fold decrease in CD34^+^CD38^−^CD7^+^ (DN1) TSPs at week 2 post infection. **b,** Correlation between frequency of peripheral T cells and ex-vivo generated T cells from CD34^+^CD3^−^ HSPCs isolated from BM of RMs at week 16 post infection. **c-f,** Correlation between fold decrease in bone marrow (DN1) TSPs at weeks 2 and plasma viral load at week 2 (c), week 4 (d), week 12 (e) and week 16 (f) post SIV infection. **g–j,** Correlation between the frequency of ex-vivo generated T cells from CD34^+^CD3^−^ HSPCs isolated from BM of RMs at week 16 post infection (showing pre-thymic T cell potential of progenitors) and plasma viral load at week 2 (g), week 4 (h), week 12 (i) and week 16 (j) post SIV infection. **k,** Correlation between total plasma viral load [AUC, Wk0-Wk16] and peripheral CD3+ T cells at week 16 post infection. Each symbol represents an individual animal (N=6). Solid red lines indicate linear regression. Pearson correlation coefficients (r) and corresponding P values are indicated in each panel. Red/black lines denote linear regression fits; (*P < 0.05).

**Supplementary Figure 4.**
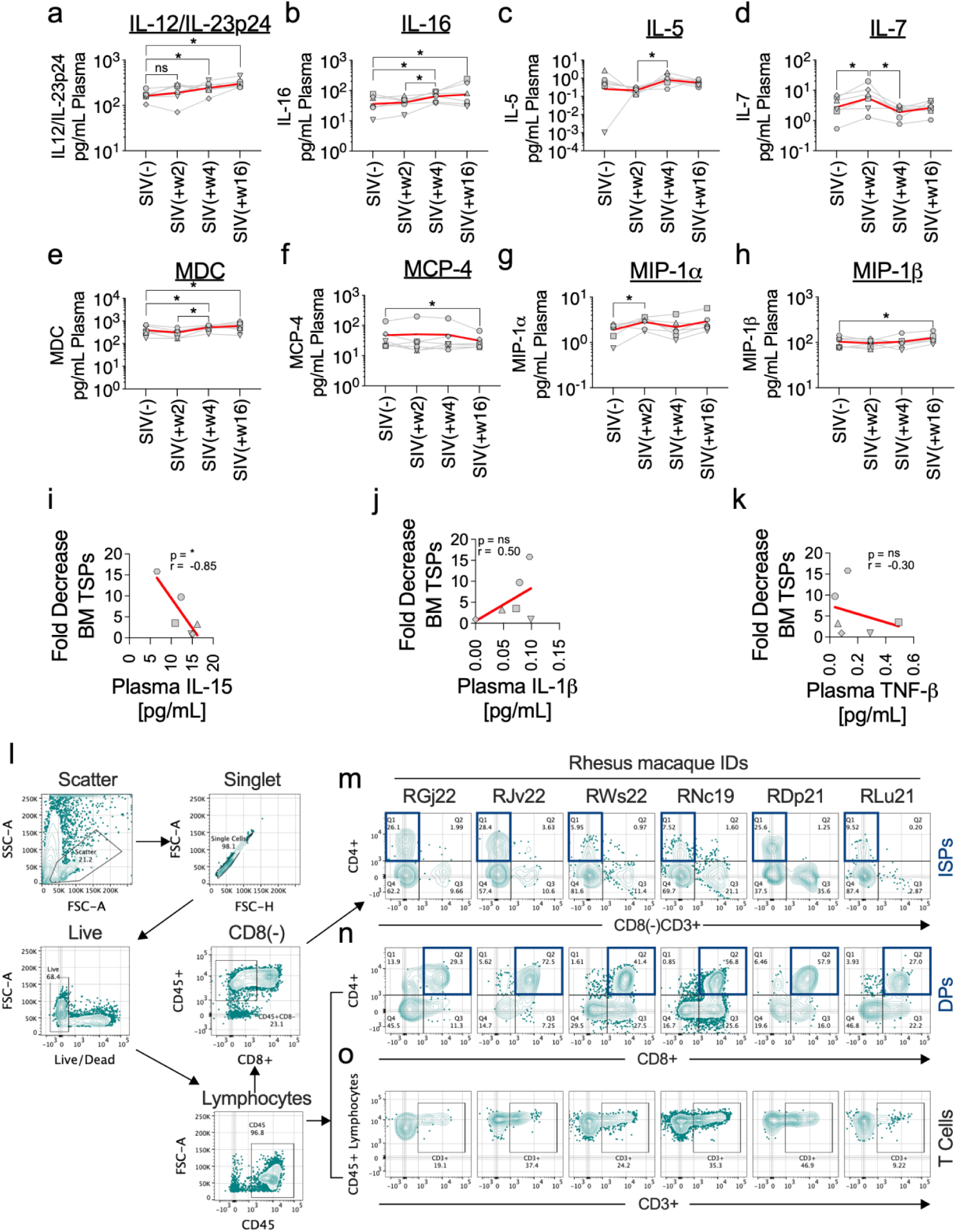
Association of plasma cytokine and chemokine signatures with T cell lineage commitment of progenitors (a–o). **a-h,** Longitudinal analysis of plasma cytokine and chemokine levels measured at baseline (SIV⁻) and at weeks 2, 4 and 16 following SIV infection, including IL-12/IL-23p24 (a), IL-16 (b), IL-5 (c), IL-7 (d), MDC (e), MCP-4 (f), MIP-1α (g) and MIP-1β (h). Data are shown as individual animals connected across time points, with group means indicated by red lines. **i–k,** Correlation between the fold decrease in bone-marrow CD34^+^CD38^−^CD7^+^ (DN1) TSPs and plasma levels of IL-15 (i), IL-1β (j) and TNF-β (k) at peak viremia (week 2 post infection). Each symbol represents an individual animal. Solid red lines indicate linear regression. Pearson correlation coefficients (r) and P values are shown, ns, not significant. **l-o,** Representative flow-cytometric plots showing gating strategy for analysis of T cell lineage committed differentiation of CD3^−^CD34⁺ HSPCs isolated from bone marrow of rhesus macaques (RMs) at week 16 post infection. Cells were gated on scatter, singlets, live, CD45⁺ cells, followed by identification of CD4⁺CD3⁻ immature thymocytes (within CD8^−^CD45^+^; m), CD45^+^ CD4⁺CD8⁺ double positive (n), as well as mature CD45^+^CD3⁺ T cells (o). Statistical significance was assessed using paired Wilcoxon t-tests. Pearson correlation coefficients (r) and corresponding P values are indicated in each panel. (*P < 0.05).

**Supplementary Figure 5.**
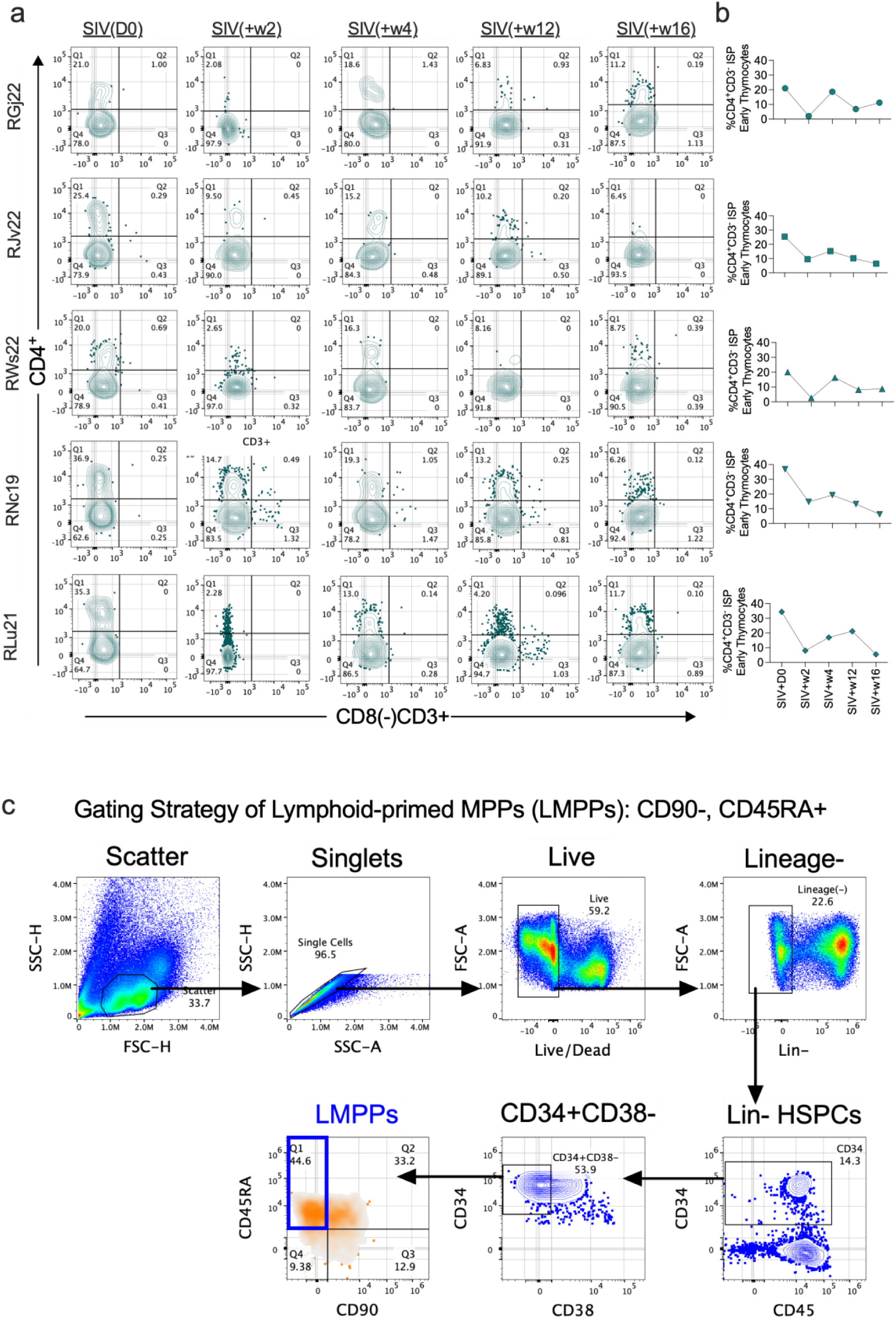
Bone marrow–derived HSPCs exhibit reduced thymocyte regenerative capacity during pathogenic SIV infection (a-c), a, Representative flow-cytometric plots showing CD4 versus CD3 expression among RhATO-derived thymocytes generated from bone marrow HSPCs collected at baseline (SIV D0) and at weeks 2, 4, 12 and 16 post- SIV infection. Data are shown for individual rhesus macaques (RGj22, RJv22, RWs22, RNc19 and RLu21). A progressive reduction in the frequency of CD4⁺CD3⁻ early immature single-positive thymocytes is observed following SIV infection. b, Longitudinal quantification of the frequency of CD4⁺CD3⁻ early immature single-positive thymocytes generated in RhATOs from bone marrow HSPCs at baseline and at the indicated time points following SIV infection. Each data point represents an individual animal, demonstrating a sustained decline in thymocyte regenerative potential of bone marrow derived HSPCs during progressive infection. c, Flow-cytometric gating strategy used to identify lymphoid-primed multipotent progenitors (LMPPs) from bone marrow based on reported strategy(31). Cells were sequentially gated on scatter, singlets, live cells and lineage-negative (Lin⁻) populations, followed by identification of CD34⁺ HSPCs. LMPPs were defined within the CD34⁺CD38⁻ compartment as CD90⁻CD45RA⁺ cells. Numbers indicate frequencies within parent gates.

**Supplementary Figure 6.**
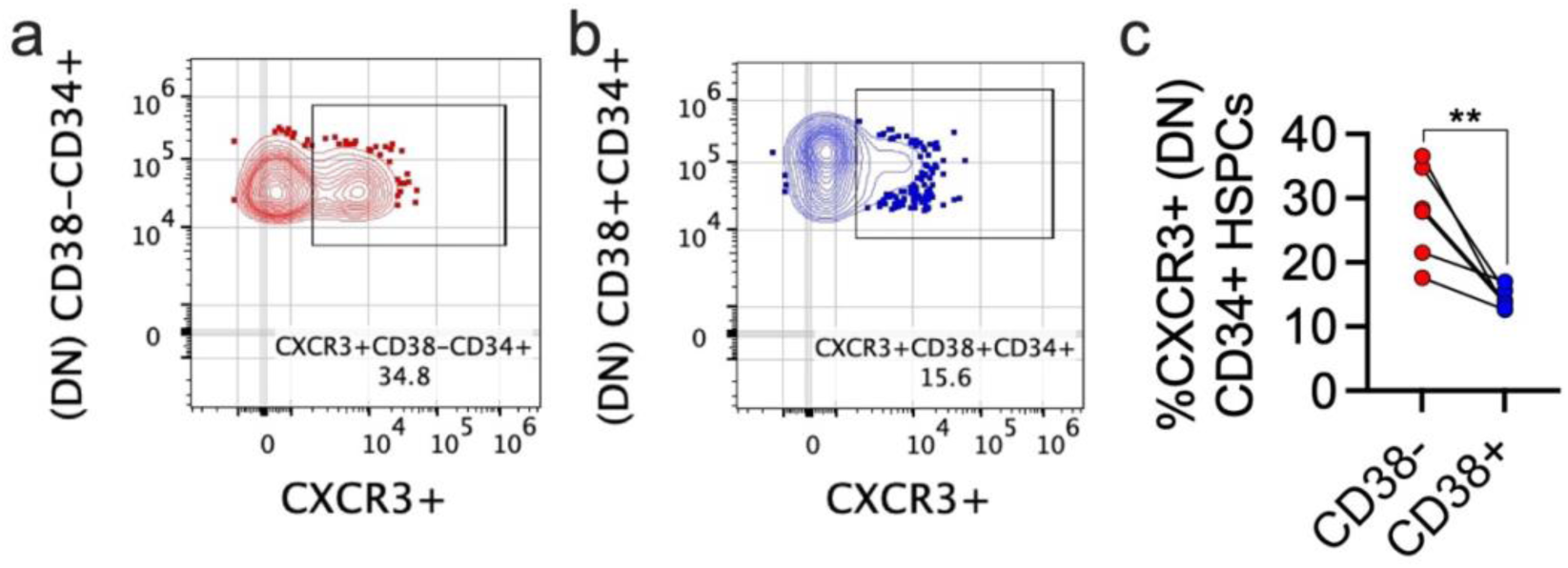
Expression of CXCR3 on CD38(-) and CD38(+) CD34+ HSPCs (a-c). **a,b,** Representative flow-cytometry plots showing CXCR3 expression on double-negative (DN; CD4⁻CD8⁻) CD34⁺ hematopoietic stem and progenitor cells (HSPCs) within the CD38⁻ (a) and CD38⁺ (b) compartments. Gates indicate CXCR3⁺ populations, with frequencies shown. c, Paired comparison of the frequency of CXCR3⁺ cells among DN CD34⁺ HSPCs within the CD38⁻ and CD38⁺ compartments. Each paired symbol represents one animal. Statistical significance was assessed using paired Wilcoxon t-tests. **P < 0.01.

**Supplementary Figure 7.**
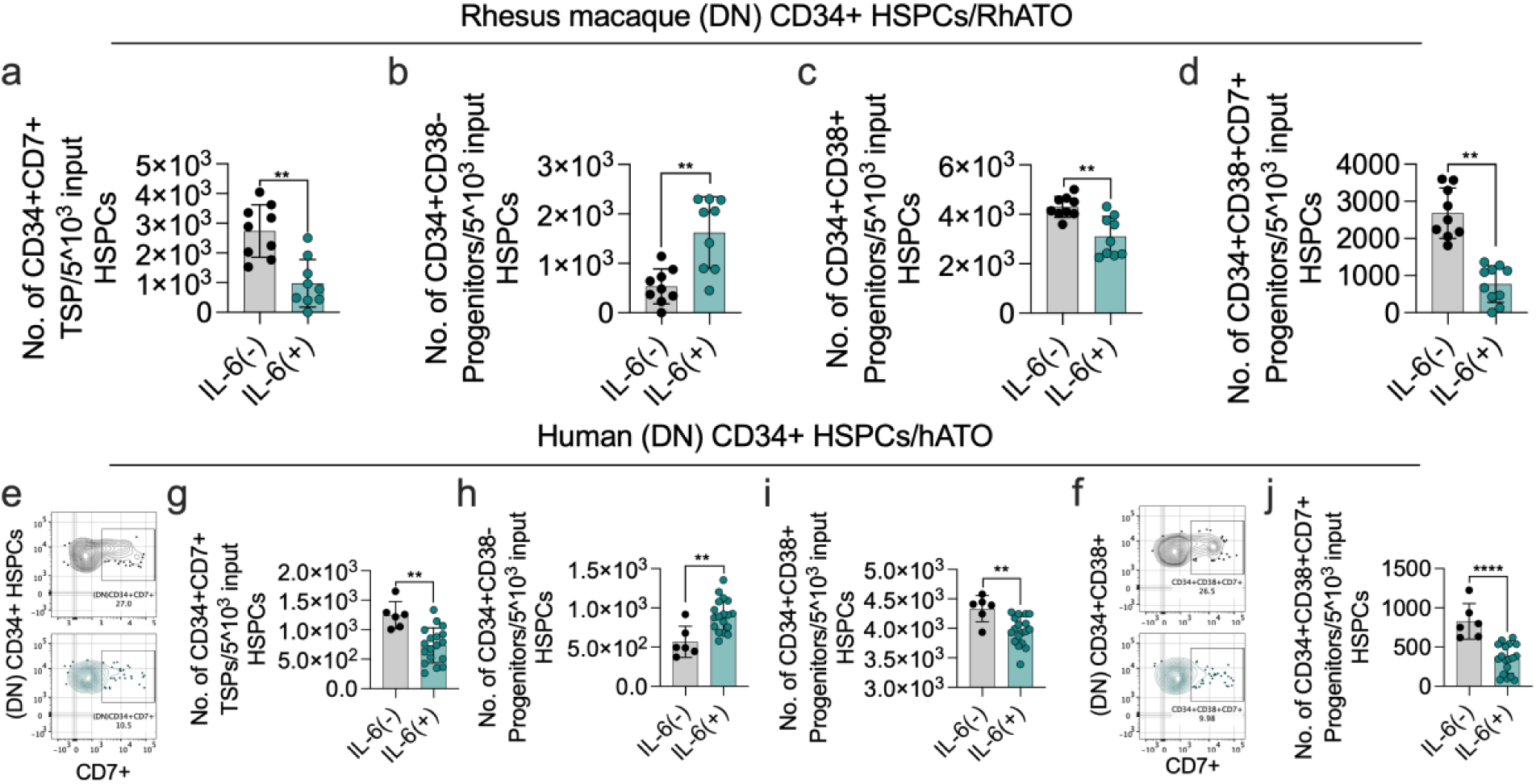
IL-6 mediated impairment of CD38 expression on CD3^−^CD34^+^ HSPCs from virus naïve bone marrow and resulting decline of CD34⁺CD38⁻CD7⁺ (DN2) population in rhesus- and human-specific nonanimal models of thymopoiesis. **a–d,** Analysis of early events of T cell lineage committed differentiation of bone marrow CD3^−^CD34^+^ HSPCs in the absence or presence of IL-6 at week 1 of RhATO culture. **a,** Bar graph showing absolute number of DN CD34⁺CD7⁺ T cell progenitors per 5 × 10³ input HSPCs generated in the absence or presence of IL-6. **b,** Bar graph showing absolute numbers of DN CD34⁺CD38⁻ progenitors per 5 × 10³ input HSPCs in the absence or presence of IL-6. **c,** Bar graph showing absolute numbers of DN CD34⁺CD38⁺ progenitors per 5 × 10³ input HSPCs in the absence or presence of IL-6. **d,** Bar graph showing absolute numbers of CD34⁺CD38⁺CD7⁺ (DN2) TSPs per 5 × 10³ input HSPCs following IL-6 treatment. **e–j,** Analysis of early events of T cell lineage committed differentiation of human bone marrow derived primary CD3^−^CD34^+^ HSPCs (HIV naïve) in the absence or presence of IL-6 at week 1 of hATO culture. **e,** Representative flow-cytometry plots showing frequency of (DN)CD34+CD7+ T cell progenitors in the absence or presence of IL-6 at week 1 of hATO culture. **g,** Bar graph showing absolute number of DN CD34⁺CD7⁺ T cell progenitors per 5 × 10³ input HSPCs generated in the absence or presence of IL-6. **h,** Bar graph showing absolute numbers of DN CD34⁺CD38⁻ progenitors per 5 × 10³ input HSPCs in the absence or presence of IL-6. **i,** Bar graph showing absolute number of DN CD34⁺CD38⁺ progenitors per 5 × 10³ input HSPCs in the absence or presence of IL-6. **j,** Bar graph showing absolute numbers of CD34⁺CD38⁺CD7⁺ (DN2) TSPs generated per 5 × 10³ input HSPCs in the absence or presence of IL-6. Each symbol represents an independent donor or biological replicate. Data are shown as geomean ± geomean s.d. Statistical significance was determined using Mann Whitney unpaired two-tailed t-tests. (**P < 0.01; **P < 0.0001).

**Supplementary Table 1.** Table showing information of six Rhesus macaques recruited in this study for longitudinal study.

| Sr. No. | Animal Code | Sex | Age | Weight (Kg) | B08 | B17 | A01 |
| --- | --- | --- | --- | --- | --- | --- | --- |
| 1 | RGj22 | M | 3y 4m | 4.71 | - | - | + |
| 2 | RJv22 | M | 2y 6m | 4.44 | - | - | + |
| 3 | RWs22 | M | 2y 7m | 4.58 | - | - | + |
| 4 | RNc19 | F | 6y 8m | 5.7 | - | - | - |
| 5 | RDp21 | F | 4y 3m | 7.25 | - | - | - |
| 6 | RLu21 | F | 3y 6m | 4.84 | - | - | - |

